# preciseTAD: A transfer learning framework for 3D domain boundary prediction at base-pair resolution

**DOI:** 10.1101/2020.09.03.282186

**Authors:** Spiro C. Stilianoudakis, Maggie A. Marshall, Mikhail G. Dozmorov

**Affiliations:** Dept. of Biostatistics, Dept. of Pathology, Virginia Commonwealth University, Richmond, VA, 23298, USA; Bioinformatics program, Virginia Commonwealth University, Richmond, VA, 23298, USA

## Abstract

Chromosome conformation capture technologies (Hi-C) revealed extensive DNA folding into discrete 3D domains, such as Topologically Associating Domains and chromatin loops. The correct binding of CTCF and cohesin at domain boundaries is integral in maintaining the proper structure and function of these 3D domains. 3D domains have been mapped at the resolutions of 1 kilobase and above. However, it has not been possible to define their boundaries at the resolution of boundary-forming proteins. To predict domain boundaries at base-pair resolution, we developed preciseTAD, an optimized transfer learning framework trained on high-resolution genome annotation data. In contrast to current TAD/loop callers, preciseTAD-predicted boundaries are strongly supported by experimental evidence. Importantly, this approach can accurately delineate boundaries in cells without Hi-C data. preciseTAD provides a powerful framework to improve our understanding of how genomic regulators are shaping the 3D structure of the genome at base-pair resolution.

## Introduction

The advent of chromosome conformation capture (3C) sequencing technologies, and its successor Hi-C, have revealed a hierarchy of the 3-dimensional (3D) structure of the human genome such as chromatin loops,^1,2^ Topologically Associating Domains (TADs),^3–5^ and A/B compartments,^2,6^ reviewed in Beagan and Phillips-Cremins^7^ and Chang et al..^8^ At the kilobase scale, chromatin loops (corner-dot structures on Hi-C chromatin interaction maps) connect gene promoters with distal enhancers and regulate gene expression. At the megabase scale, TADs represent regions on the linear genome that are highly self-interacting. Perhaps the most prominent feature of TADs is that they are demarcated by boundaries constraining enhancer-promoter interactions,^9^ although these constraints can be flexible.^10^ Perturbation of boundaries have been reported to promote human cancers,^11,12^ neurological disorders,^13^ and pathologies of limb development.^1,14^ Identifying the precise location of boundaries remains a top priority to fully understand the functionality of the human genome.

Several methods have been proposed to identify TAD boundaries (reviewed in),^15^ and chromatin loops.^2,16,17^ They are primarily based on identifying characteristic patterns in Hi-C contact matrices, such as dense inter-TAD contacts, sparse intra-TAD contacts, among other features.^4,7^ Consequently, they are limited by Hi-C data resolution. Resolution refers to the size of genomic regions (bins) used to segment the linear genome and create Hi-C contact matrices. Lower resolution corresponding to larger bin sizes leads to increased uncertainty in domain boundary location.

TAD and loop boundaries are not mutually exclusive. Rao et al. demonstrated that boundaries of TADs, referred to as “contact domains”, are enriched in chromatin loops.^2^ Numerous observations demonstrated the presence of hierarchically nested chromatin domains within TADs.^2,18^ Domain boundaries are thought to form by the ‘loop extrusion’ mechanism. During extrusion, the molecular motors (condensin and cohesin) track along the DNA sequence ‘extruding’ the intervening DNA in an ATP-dependent manner and pausing at the convergent CTCF motifs.^19–25^ Consequently, boundaries are expected to be enriched in CTCF and RAD21/SMC3, members of the cohesin complex.^2,4,18,26,27^ More recently, ZNF143 has been identified as a cofactor of CTCF-Cohesin complex.^28,29^ Furthermore, distinct patterns of histone modifications have also been shown to be present at boundaries.^4,6^

In contrast to low-resolution Hi-C matrices, functional/regulatory genomic annotations (histone modifications, DNAse I hypersensitive sites, DNA methylation, and transcription factor binding sites (TFBSs)) have been profiled at a relatively high resolution (10-300bp).^30,31^ Genomic annotations have been used to predict functional chromatin contacts (e.g., HiC-Reg),^32^ boundaries of chromatin domains (e.g., nTDP,^33^ Lollipop,^34^ 3DEpiLoop,^35^ TAD-Lactuca),^36^ with 48 methods recently reviewed by Tao et al.^37^ (see also Belokopytova & Fishman 2020 for a broader overview).^38^ Yet, these methods operate at the resolution of Hi-C data. Because increasing resolution of Hi-C data requires a quadratic increase in sequencing depth^39^ and the associated costs, most currently available Hi-C matrices have relatively low resolution, ranging from 1 kb to 100 kb. Furthermore, conventional Hi-C relies on a 0.1-10 kb size fragmentation by a restriction enzyme, but the existence of self-ligated products and undigested fragments with sizes 1-10 kb prohibits analysis at higher resolution.^2,40^ The association of domain boundaries with genomic annotations suggests these annotations may inform the more precise location of domain boundaries.

We present *preciseTAD*, an optimally tuned transfer learning framework for precise domain boundary detection using genomic annotation data. *preciseTAD* learns the associations between genomic annotations and boundaries detected from low-resolution Hi-C matrices and transfers the learned associations at base-level resolution (predicts the probability of each base being a boundary). This approach circumvents resolution restrictions of Hi-C matrices and allows for the precise detection of domain boundaries. We demonstrate that *preciseTAD*-predicted boundaries are strongly enriched in known molecular drivers of 3D chromatin including CTCF, RAD21, SMC3, and ZNF143. Further, we show that the associations learned in one cell line can be used to predict boundaries in other cell lines using cell-specific genomic annotations only. As such, *preciseTAD* allows for predicting domain boundaries in cells without Hi-C data. We provide domain boundary predictions for 60 cell lines, demonstrating that imputing missing cell-specific genomic annotations with Avocado^41^ is a viable approach to recover domain boundaries. The *preciseTAD* R package and the pre-trained models (*preciseTADhub* ExperimentHub package) are freely available on Bioconductor.

## Results

### preciseTAD overview

*preciseTAD* implements the idea of transfer learning across genomic data resolutions. It models the association between chromatin domain boundaries and genomic annotations using low-resolution (5-100kb) Hi-C genomic regions and applies this model at base-level resolution to predict the probability of each base being a boundary (Figure 1). Our method utilizes the random forest (RF) algorithm trained on chromatin state (BroadHMM), histone modification (HM), and transcription factor binding site (TFBS) annotation data. Our training/testing framework was used to determine the optimal set of data-level characteristics including resolution (bin size), feature engineering, and resampling (Figure S1). We found that spatial associations (linear distance) between boundaries and genomic annotations perform best, transcription factor binding sites outperform other annotations, and a simple random undersampling technique addresses the negative effect of class imbalance. *preciseTAD* employs density-based clustering (DBSCAN) and partitioning around medoids (PAM) to detect annotation-guided boundary regions and summit points with the highest boundary probability. These improved domain boundary locations can provide insight into the association between genomic regulators and the 3D genome organization.

**Figure 1.**
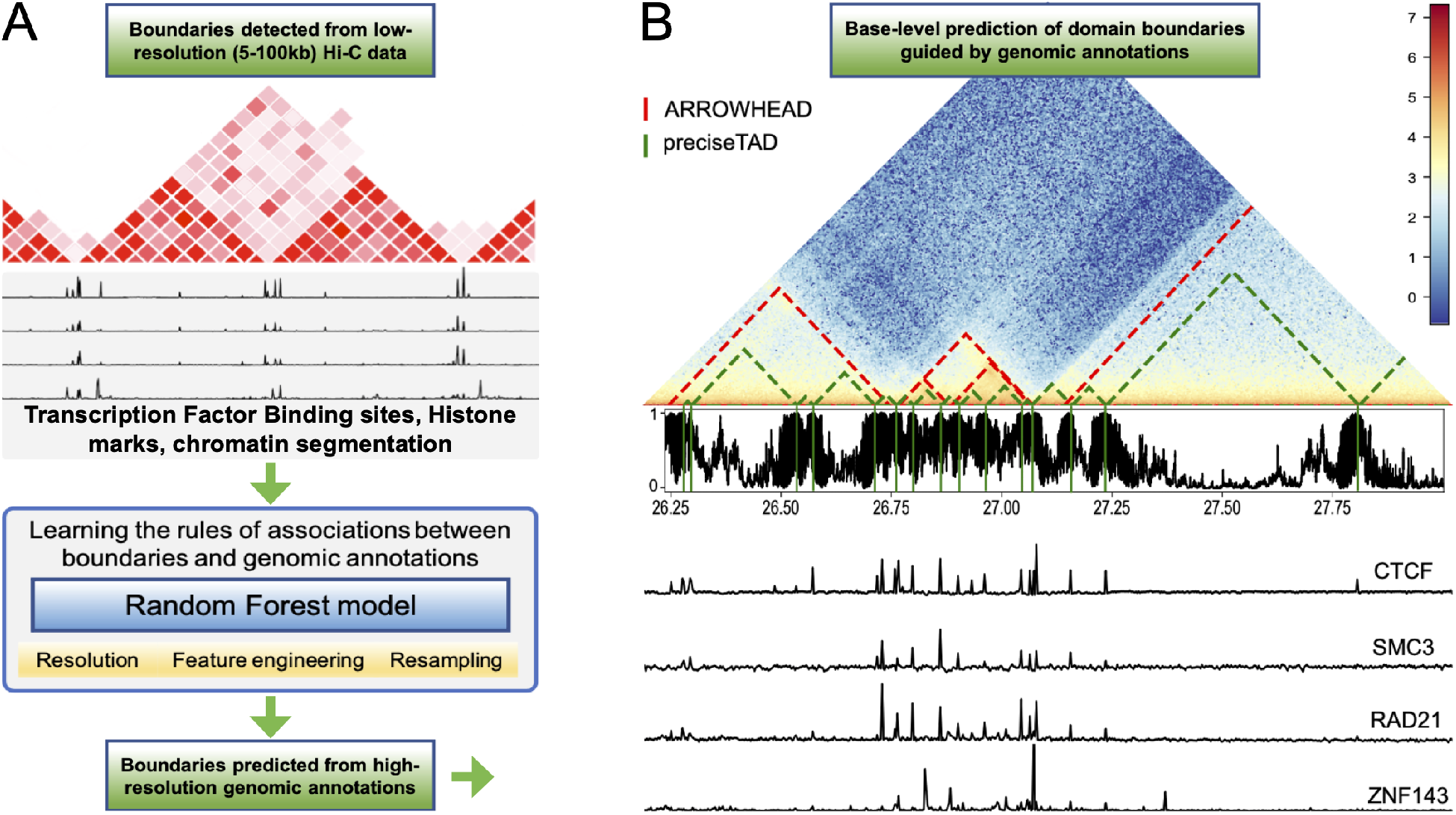
Overview of preciseTAD. (A) The framework of preciseTAD includes Random Forest models learning the optimal rules of associations between low-resolution boundaries and genomic annotations, and predicting the probability of each base being a boundary. First, feature engineering steps are applied to capture various rules of boundary-annotation associations. Second, the models are trained on different data resolution, addressing class imbalance. Third, the most predictive features shared across chromosomes and cell lines are selected for the final model. Fourth, a base-level predictor matrix is build, in which each base is annotated with the most optimal features and association rules. Fifth, the probability of each base being a boundary is predicted with the optimal model trained on low-resolution Hi-C data. (B) The final step includes clustering bases with high boundary probability into preciseTAD boundary regions (PTBRs) using DBSCAN and identifying the most likely boundary points (PTBPs) using PAM. The output of preciseTAD includes genomic coordinates of PTBRs, PTBPs, and their summary statistics.

### Developing an ML framework for optimal boundary prediction

We developed a machine learning (ML) framework for determining the optimal set of data level characteristics to predict boundary regions of Topologically Associating Domains (TADs) and chromatin loops, collectively referred to as domain boundaries. Similar to other boundary prediction methods,^17,32,34,35,42^ we chose the random forest (RF) algorithm as our binary classification tool. The reason for it is two-fold: (1) to devise a tunable prediction rule in a supervised learning framework that is both robust to overfitting and able to handle multiple correlated predictors, and (2) to allow for an interpretable ranking of predictors.^43^ Furthermore, previous reports demonstrated robustness of RF to overfitting and superior performance over other machine learning classifiers.^35,36,42^

As an example of TAD boundaries, we derived cell line-specific boundaries at 5-100 kb resolutions using *Arrowhead*.^2^ For chromatin loops, we used loops derived by *Peakachu*^17^ (Table S1). We chose GM12878 (lympho-blastoid) and K562 (chronic myelogenous leukemia) as cell lines with the most rich and comparable sets of genomic annotations. The choice of *Peakachu* loops was motivated by their good overlap with HiCCUP^2^ and Fit-Hi-C^16^ loops and better enrichment in known boundary factors. To ensure the cell line-specific predictions corresponded to experimental domain boundaries, we used cohesin-bound chromatin loop data from ENCODE phase 3,^44^ referred hereafter as *Grubert* boundaries (Table S1). We found that boundaries Arrowhead/Peakachu boundaries showed markedly little enrichment in binding of CTCF and other architectural proteins as compared with Grubert boundaries (Figure S10AC), further motivating the need for more precise boundary detection.

The total number of called TADs, their unique boundaries, and the number of genomic bins expectedly decreased with the decreased resolution of Hi-C data (Table 1, Table S2). The number of non-boundary regions highly outnumbered boundary regions. Such a disproportional presence of examples in one class is known as a “class imbalance” problem that negatively affects predictive modeling.^45^ To address class imbalance, we evaluated the effect of three resampling techniques. *Random over-sampling* (ROS) was defined as sampling with replacement from the minority class (boundary regions). *Random under-sampling* (RUS) was defined as sampling with replacement from the majority class (non-boundary regions). Lastly, we tested *Synthetic minority over-sampling technique* (SMOTE), which is a combination of both random over- and under-sampling to create balanced classes^46^ (see Methods).

**Table 1.** Domain boundary data and class imbalance summaries across resolutions for Arrowhead, Peakachu, and Grubert data in GM12878 cell line.

| Tool | Resolution/Bin Size | Total number of called TADs/chromatin loops | Total number of unique TAD/chromatin loop boundaries | Total number of genomic bins | Class imbalance |
| --- | --- | --- | --- | --- | --- |
| Arrowhead | 5 kb | 8052 | 15468 | 535363 | 0.03 |
| Arrowhead | 10 kb | 7676 | 14253 | 267682 | 0.05 |
| Arrowhead | 25 kb | 4670 | 8363 | 107073 | 0.08 |
| Arrowhead | 50 kb | 2349 | 4224 | 53537 | 0.08 |
| Arrowhead | 100 kb | 1031 | 1883 | 26768 | 0.07 |
| Peakachu | 10 kb | 16185 | 21421 | 267682 | 0.14 |
| Grubert | 5 kb | 16232 | 18455 | 535363 | 0.07 |

Our models were trained on cell line-specific functional genomic annotation data from ENCODE.^31^ A total of 77 cell line-specific genomic annotations were used to build the predictor space. These included histone modification (HM) data previously shown to be useful for boundary predictions,^33,35,36^ Broad ChromHMM chromatin segmentation data that captures regions with similar epigenetic activity patterns (BroadHMM), and transcription factor binding sites (TFBS, Table S3). Boundary regions were defined as genomic bins containing a called boundary (Y = 1), while non-boundary regions were defined as bins that did not contain a called boundary (Y = 0, Figure S2, see Methods). Four feature engineering procedures were developed to quantify the association between genomic annotations and bins. These included signal strength association (Signal), direct (OC), proportional (OP), and spatial (*log*_2_ + 1 Distance) relationships (Figure S1, see Methods). In total, we considered combinations of data from two cell lines *L = {GM12878, K562}*, five resolutions *R = {5 kb, 10 kb, 25 kb, 50 kb, 100 kb}*, four types of predictor spaces *P = {Signal, OC, OP, Distance}*, and three re-sampling techniques *S = {None, RUS, ROS, SMOTE}*. Once the model inputs were established, a random forest classifier was trained on *n* – 1 autosomal chromosomes, while reserving the *i^th^* chromosome for testing. Three-fold cross-validation was used to tune the *mtry* hyperparameter, while *ntree* and *nodesize* were fixed at 500 and at 0.1% of the rows in the training data, respectively. Models were validated on the testing data using cell line-specific Grubert-defined boundaries as **Y***_test_*. Model performance was evaluated by aggregating the mean balanced accuracy (BA) across each holdout chromosome, with additional performance metrics (accuracy, AUROC, AUPRC) shown in Table S4. These strategies allowed us to select the best performing model characteristics (Figure S1, see Methods).

### Random under-sampling, distance-based predictors, and high-resolution Hi-C data provide optimal performance for boundary prediction

When using Arrowhead data with class imbalance, the models exhibited low balanced accuracies, with minimal variability among different resolutions (Figure 2). Similarly, poor performances were found when using ROS. However, RUS and SMOTE re-sampling led to a drastic improvement in performance, especially at higher resolutions. We found that RUS marginally outperformed SMOTE under most conditions and used it for the subsequent analyses unless noted otherwise.

**Figure 2.**
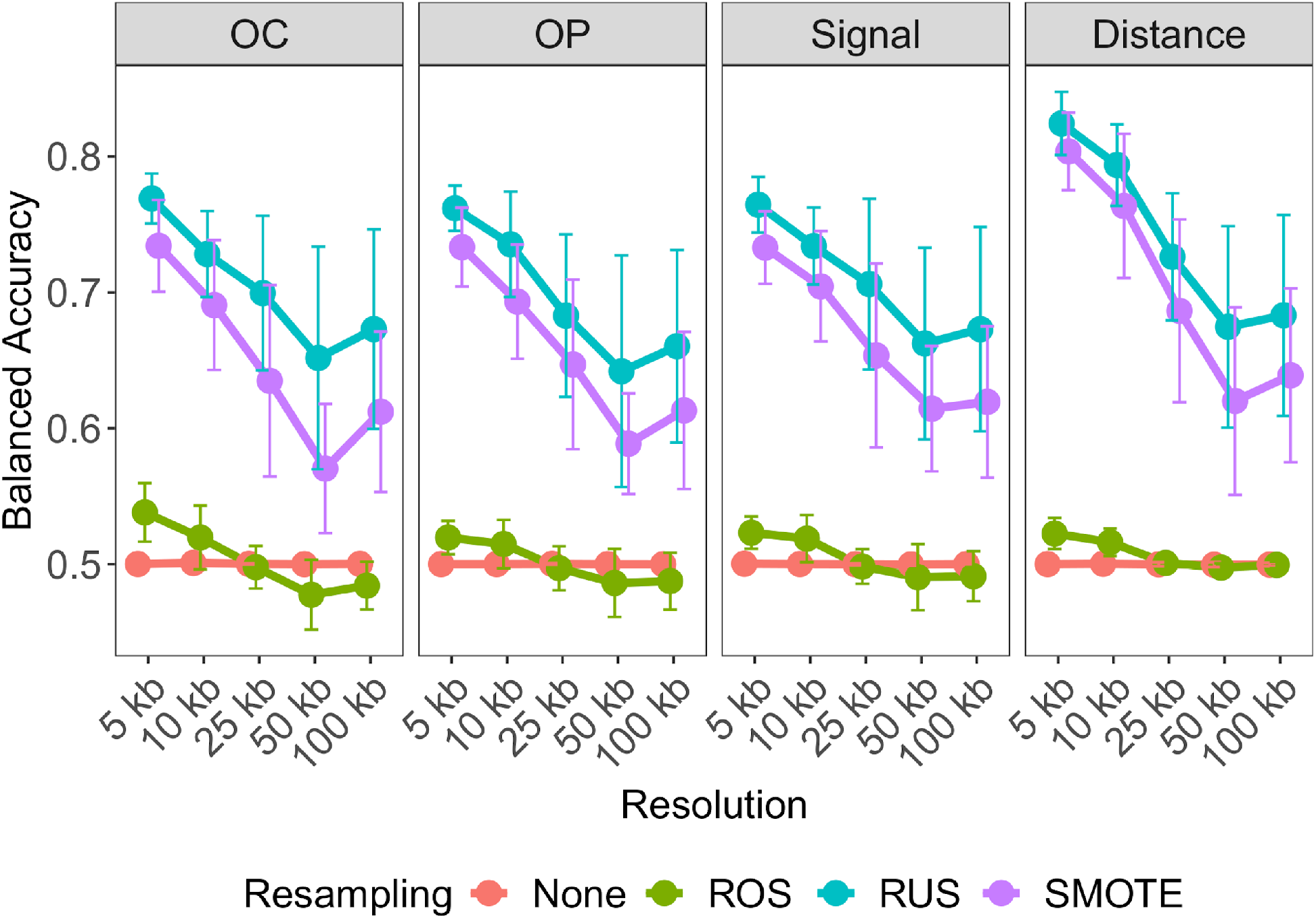
Determining optimal data level characteristics for building TAD boundary region prediction models on GM12878. Averaged balanced accuracies are compared across resolution, within each predictor-type: overlap count (OC), overlap percent (OP), average Signal and Distance, and across resampling techniques: no resampling (None; red), random over-sampling (ROS; green), random under-sampling (RUS; blue), and synthetic minority over-sampling (SMOTE; purple). Error bars indicate 1 standard deviation from the mean performance across each holdout chromosome used for testing.

Additionally, we found that distance-type predictor space yielded substantially higher balanced accuracy than the peak signal strength, overlap count, and overlap percent predictor types. As with class balancing techniques, this improvement was less evident at lower resolutions, with results consistent for K562 (Figure S4A). Furthermore, 5 kb resolution genomic bins led to the optimal prediction for TAD boundary regions on both cell lines. These observations were replicated when Peakachu-defined loop boundary regions were used (Figure S4BC). Our results indicate that random under-sampling, distance-type predictors, and high-resolution Hi-C data provide the optimal set of data characteristics for both TAD and loop boundary prediction.

### Transcription factor binding sites outperform histone- and chromatin state-specific models

We hypothesized that the class of genomic annotation may also affect predictive performance. Using the established optimal settings (RUS, Distance, 5kb/10kb resolution), we used histone modifications (HM), chromatin states (BroadHMM), and transcription factor binding sites (TFBS) to build the predictor space. Despite previous success in using histone modifications for TAD boundary predictions,^32,35,36^ their performance in our settings was least optimal. BroadHMM segmentations also performed less optimally, in agreement with previous observations.^33^ We found that TFBSs outperformed other annotation-specific models, with results consistent for loop boundaries, on both cell lines (Figure 3A; Figure S5A), and the use of all genomic annotations did not significantly improved model performance. These results suggest that transcription factors are the primary drivers of Arrowhead/Peakachu-defined boundaries in both GM12878 and K562 cell lines.

**Figure 3.**
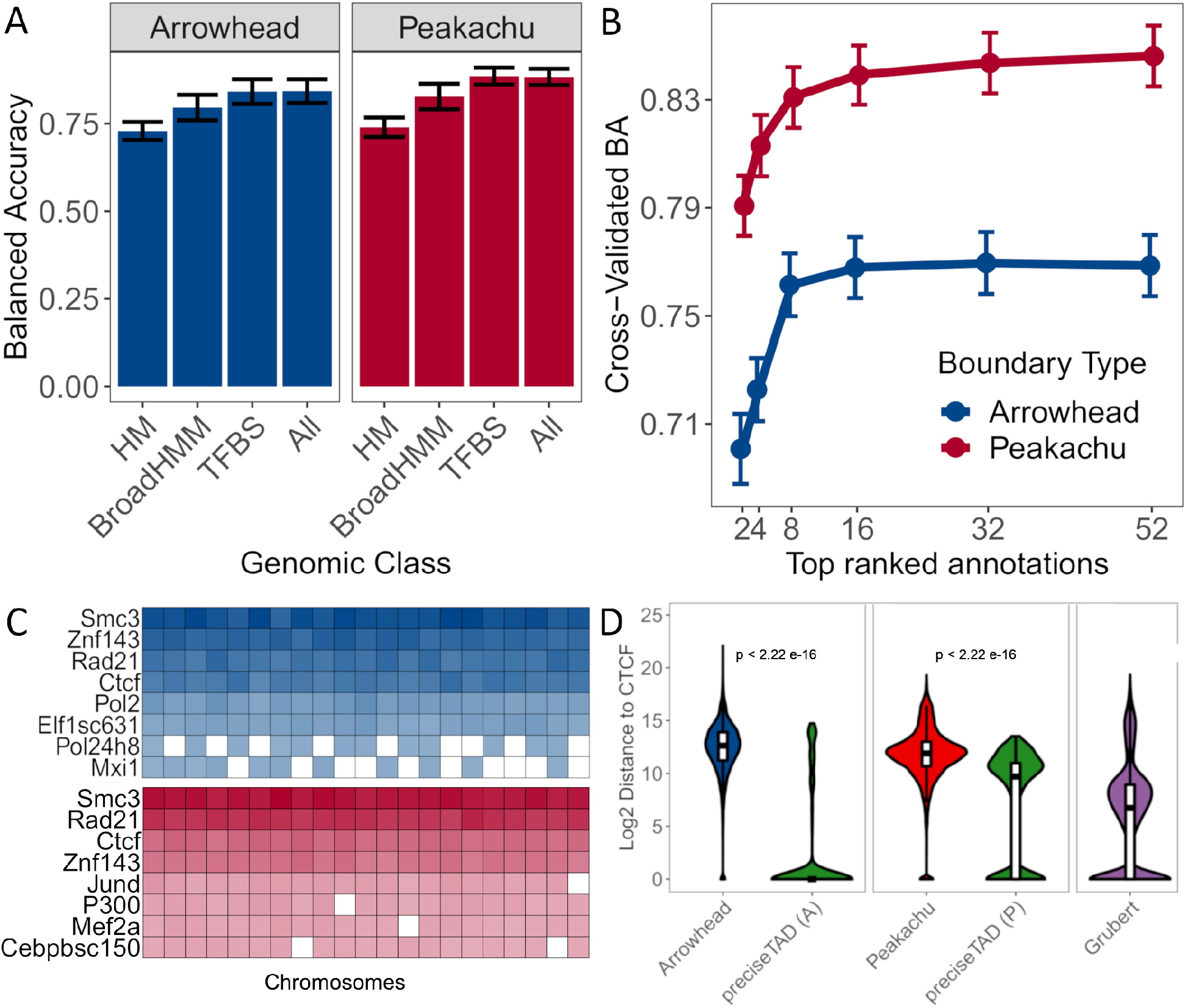
SMC3, RAD21, CTCF, and ZNF143 transcription factors accurately predict TAD and loop boundaries in GM12878. (A) Barplots comparing performances of TAD (Arrowhead) and loop (Peakachu) boundary prediction models using histone modifications (HM), chromatin states (BroadHMM), transcription factor binding sites (TFBS), in addition to a model containing all three classes (ALL). (B) Recursive feature elimination (RFE) analysis used to select the optimal number of predictors. Error bars represent 1 standard deviation from the mean cross-validated accuracy across each holdout chromosome. (C) Clustered heatmap of the predictive importance for the union of the top 8 most predictive chromosome-specific TFBSs. The columns represent the holdout chromosome excluded from the training data. Rows are sorted in decreasing order according to the columnwise average importance. (D) Violin plots illustrating the *log*_2_ genomic distance distribution from original Arrowhead/Peakachu boundaries vs. *preciseTAD*-predicted boundaries to the nearest CTCF sites. The p-values are from the Wilcoxon Rank Sum test.

### Predictive importances confirmed the biological role of CTCF, RAD21, SMC3, and ZNF143 for boundary formation

We sought to further optimize our boundary region prediction models. We implemented recursive feature elimination to avoid overfitting and selected only the most influential features across all chromosomes. We were able to obtain near-optimal performance using approximately eight TFBS (Figure 3B; Figure S5B). However, given that we trained our models on chromosome-specific data, the most significant annotations varied for each chromosome. To determine transcription factors most important for genome-wide boundary prediction, we clustered the predictive importance (mean decrease in accuracy) of the top eight significant TFs across chromosomes. We found four transcription factors, CTCF, RAD21, SMC3, and ZNF143, being consistently predictive of TAD and loop boundaries (Figure 3C; Figure S5C). Although CTCF and cohesin binding are known to colocalize, cohesin peaks are slightly shifted to the 3’ ends of convergently oriented motifs^21,27^ and appear to complement the model’s performance. We selected these top four TFBS when building the random forest model, thereby decreasing computational burden while maintaining high predictive power. Comparison of distance-to-nearest-CTCF distribution between original Arrowhead/Peakachu boundaries and *preciseTAD*-predicted boundaries showed that the latter were located closer to CTCF binding sites (Figure 3D, Figure S5D), and these results were observed for other transcription factors (Figure S6). In summary, our model was able to yield the known molecular drivers of the loop extrusion model,^19–24^ suggesting that TAD and loop boundary formation may be carried out by similar mechanisms.^7^

### *preciseTAD* identifies precise and biologically relevant domain boundaries

Using our optimally built random forest model trained on Arrowhead/Peakachu boundaries, we attempted to predict the more precise location of boundaries at base-level resolution. Intuitively, instead of bin-level annotations, the predictor-response space was built on a base-level. That is, each base was annotated with the distance to the nearest/overlapping CTCF, RAD21, SMC3, and ZNF143 site. The model trained on a bin-level space was then applied on a base-level space to predict each base’s probability of being a boundary. Our method, referred to as *preciseTAD*, uses density-based spatial clustering (DBSCAN) and partitioning around medoids (PAM) to cluster bases with high probability of being a boundary into boundary regions (PTBRs) and summit points (PTBPs, see Methods; Figure S7). We found that Arrowhead PTBRs and Peakachu PTBRs were highly overlapping (Jaccard 0.606/0.757, GM12878/K562 cell line, respectively) as compared with the less overlapping original Arrowhead/Peakachu boundaries (Jaccard 0.227/0.199, Figure S8A). Similarly, Arrowhead PTBRs and Peakachu PTBRs showed better agreement with experimental Grubert data (e.g., original Arrowhead-Grubert Jaccard 0.260 vs. Arrowhead PTBRs-Grubert Jaccard 0.292, GM12878 cell line), and there results were consistent in K562 cell line (Figure S8B). These results suggest that *preciseTAD* identifies similar boundaries when trained on either Arrowhead or Peakachu data, and these boundaries better agree with experimentally observed data.

When trained using Arrowhead and Peakachu boundaries at 5 kb and 10 kb, respectively, the *preciseTAD* model predicted a total of 12,258 domain and 15,707 chromatin loop boundaries in GM12878, as well as 9,603 TAD and 11,154 chromatin loop boundaries in K562 cell line (Table S5). To evaluate the biological significance of *preciseTAD* PTBRs, we investigated signal distribution of four known molecular drivers of 3D chromatin (CTCF, RAD21, SMC3, and ZNF143) around boundaries detected by different methods. *preciseTAD*-predicted boundary points (the base-level boundary locations, PTBPs) showed much stronger signal distribution than Arrowhead and Peakachu boundaries, and frequently outperformed experimentally obtained Grubert data (Figure 4AB, Figure S9AB). Notably, boundaries called by other callers similarly lacked signal distribution specificity (Figure S10AC). Surprised by the poor performance of domain boundaries detected from Hi-C data, we compared signal distribution around *preciseTAD* PTBPs with boundaries reported by Lollipop, a method for domain boundary prediction from genome annotation data and various domain characteristics.^34^ Notably, *preciseTAD*- and Lollipop-predicted boundaries showed comparable signal distribution, cementing the importance of genomic annotations for domain boundary prediction (Figure S10BD). Our results indicate that *preciseTAD*-predicted boundaries (PTBRs) and boundary points (PTBPs) better reflect the known biology of boundary formation when compared to boundaries called solely from Hi-C contact matrices.

**Figure 4.**
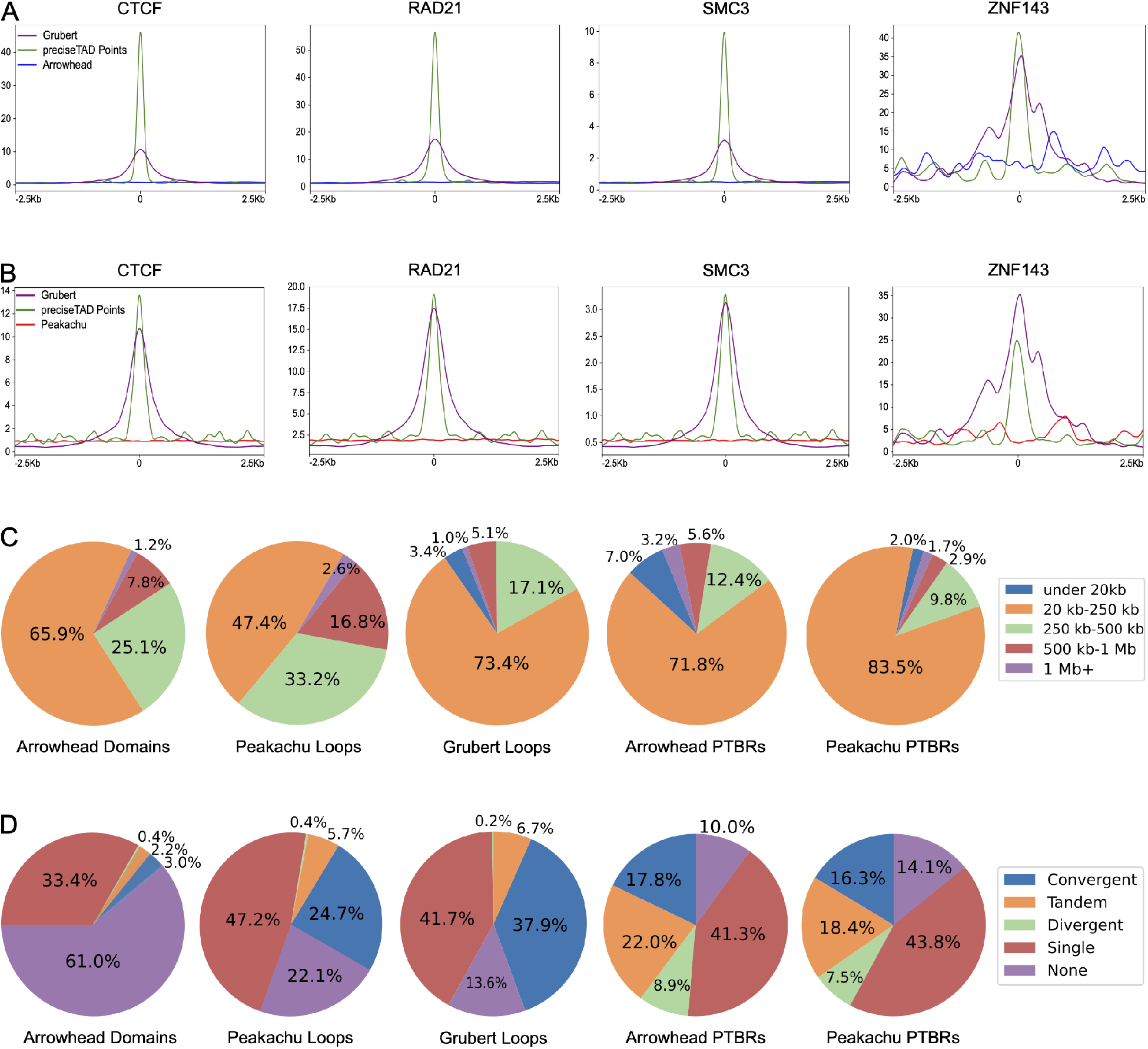
*preciseTAD* boundaries are more enriched for known molecular drivers of 3D chromatin. Signal enrichment strength of CTCF, RAD21, SMC3, and ZNF143 sites around midpoints of *preciseTAD*-predicted boundaries (green) compared to midpoints of (A) Arrowhead-called boundaries (blue), (B) Peakachu loop boundaries (red). Data for midpoints of Grubert cohesin loop boundaries is shown as a proxy for experimental “ground truth” (purple). Panel inserts show signal enrichment around preciseTAD boundary points vs. Grubert ground truth. (C) Domain size distribution, and (D) CTCF orientation analysis. Data for GM12878 cell line is shown.

We further investigated domain size distribution (Figure 4C, Figure S9C). We used loop size distribution in experimentally observed Grubert data as “ground truth”. We found that the original Arrowhead and Peakachu algorithms detected higher proportion of large domains (e.g., 25.1%/33.2% of 250kb-500kb domains vs. 17.1% in Grubert, Figure 4C, GM12878 data). In contrast, size distribution of domains marked by *preciseTAD*-detected PTBRs was similar to that of Grubert. When trained on either Arrowhead/Peakachu data, *preciseTAD* identified 71.8%/83.5% 20kb-250kb small-sized domains vs. 73.4% Grubert, 12.4%/9.8% 250kb-500kb mid-sized domains vs. 17.1% Grubert, and 5.6%/2.9% large-sized domains vs. 5.1% Grubert, with those results consistent in the K562 cell line (Figure 4C, Figure S9C). Importantly, the larger proportions of 20kb-250kb small-sized domains identified by *preciseTAD* are in better agreement with 80-120kb domain size estimated by microscopy and via modeling of Hi-C data.^25,47^ We should note that, in contrast to Arrowhead/Peakachu, *preciseTAD* identifies individual boundaries; consequently, we measured domain size as the distance between consecutive boundaries. Thus, we expect some excessively large domain sizes (e.g., if two PTBRs span centromere region) and very small domain sizes (e.g., PTBRs separated by gaps in poorly organized regions). Despite this limitation, our results suggest that *preciseTAD* identifies boundaries better reflecting experimentally observed domain size distribution.

Directionality of CTCF binding, e.g., convergent orientation of CTCF motifs, is a known defining feature of domain formation.^2,27^ We used CTCF orientation distribution in experimentally observed Grubert boundaries as “ground truth”. Indeed, Grubert loops contained the largest proportion of convergent CTCF motifs (37.9%) as compared with Arrowhead domains (3.0%). Arrowhead data contained a large propoprtion of domains lacking CTCF motifs (61.0%) or domains having single CTCF motifs (33.4%, Figure 4D). In contrast, Peakachu and *preciseTAD*-called domains showed high resemblance to CTCF motif orientation observed in Grubert data. Domains defined by the original Peakachu algorithm, Arrowhead PTBRs and Pekachu PTBRs (*preciseTAD*-defined regions trained on Arrowhead/Peakachu data) contained a high proportion of convergent CTCF motifs (24.7%, 17.8%, 16.3%), tandem motifs (5.7%, 22.0%, 18.4% vs. 6.7% Grubert), and single CTCF motifs (47.2%, 41.3%, 43.8% vs. 41.7% Grubert, Figure 4D). Domains defined by *preciseTAD* PTBRs contained the largest proportion of divergent CTCF sites (8.9% for Arrowhead PTBRs and 7.5 for Peakachu PTBRs, Figure 4D). While this may be attributed to the aforementioned non-contiguous nature of domains defined by PTBRs, these findings may reflect recent observations that domain boundaries contain divergent CTCF motifs while convergent motifs mark the interior of domains at 5-100kb range.^48^ Together with strong signal enrichment and domain size distribution results, our observations indicate that *preciseTAD* may identify domain boundaries and boundary points better reflecting known biology of boundary formation.

### Training in one cell line accurately predicts boundary regions in other cell lines

Previous studies suggest that TAD boundaries are relatively similar across cell lines.^3–5^ To assess the level of cross-cell-line similarity, we evaluated the overlap between cell line-specific boundaries detected by Arrowhead and Peakachu methods as well as *precise-TAD*-predicted boundaries trained on Arrowhead and Peakachu data. Only 24% and 30% of boundaries were overlapping between cell lines for Arrowhead and Peakachu boundaries (J=0.246 and J=0.295), respectively (Figure 5A). In contrast, *preciseTAD*-predicted boundaries were more simila between GM12878 and K562 cell lines regardless of which data was used for training (Arrowhead PTBRs overlap J=0.383; Peakachu PTBRs J=0.467, Figure 5B). This better agreement between cell type-specific *preciseTAD*-predicted boundaries further supports the notion of their higher biological relevance.

**Figure 5.**
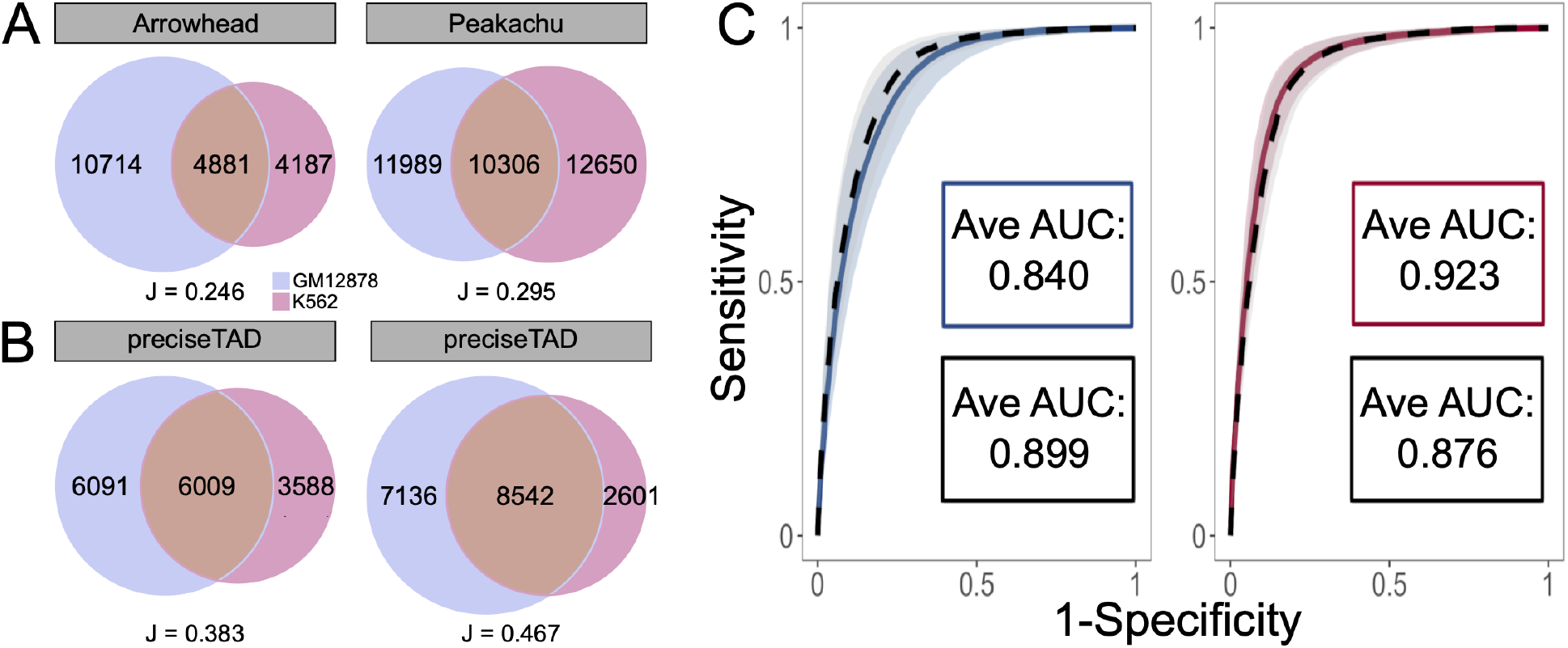
*preciseTAD* models trained in one cell line can accurately predict boundaries in another cell line. (A) Venn diagrams of overlap original Arrowhead/Peakachu domain boundaries and (B) *peakachu*-predicted PTBRs for GM12878 (red) and K562 (blue) cell lines. All boundaries were flanked by 5kb. (C) Receiver operating characteristic (ROC) curves and the corresponding average area under the curves (AUCs) when training and testing on GM12878 data (blue, Arrowhead; red, Peakachu) versus training on K562 and testing on GM12878 data (black, dashed). The curves represent the average sensitivities and specificities across each holdout chromosome. The shaded areas around each curve represent 1 standard deviation from the average.

Our observation that *preciseTAD* predicts similar domain boundaries when trained on either Arrowhead or Peakachu data raises the possibility that boundary-annotation associations learned in one cell line can predict boundaries in another cell line using its genomic annotation data. That is, given that genomic annotations (distance to CTCF, RAD21, SMC3, and ZNF143) are predictive of boundaries in one cell line, their locations in another cell line may be predictive of boundaries in that cell line. Indeed, training and testing using Arrowhead boundaries and genomic annotation data from the GM12878 cell line resulted in an average AUC=0.840 (Figure 5C). When training on the K562 boundaries/annotations and testing on GM12878, the average AUC was 0.899. Likewise, training and testing using Peakachu boundaries and genomic annotation data from the GM12878 cell line was comparable to models trained on K562 boundares/annotations and testing on GM12878 cell line (Avg. AUC=0.923 and 0.876, respectively, Figure 5A). These results were consistent when comparing training/testing strategies on K562 boundaries/annotations with training on GM12878 and testing on K562 data (Figure S11). In both instances the average ROC curves were found to be within 1 standard deviation of each other, suggesting that a model trained on data from one cell line performs well when using the data from another cell line. This ability of boundary-annotation associations learned from one cell type to successfully predict boundaries in another indicates that the same underlying forces may drive boundary formation across various cell lines.

We further evaluated biological characteristics of cell type-specific boundaries predicted by models trained on a different cell line. We evaluated two scenarios: 1) training on GM12878 and predicting boundaries on GM12878 (GM on GM) vs. training on K562 and predicting on GM12878 (K on GM), and 2) training on K562 and predicting boundaries on K562 (K on K) vs. training on GM12878 and predicting boundaries on K562 (GM on K). Using Arrowhead-trained models, 76% (J=0.701) and 81% (J=0.751) of predicted boundaries overlapped in both cross-cell-line prediction scenarios (Figure S12). When using Peakachu-trained models, we observed 85% (J=0.705) and 88% (J=0.759) overlap (Figure S13). Furthermore, boundaries predicted on unseen annotation data exhibited a similar level of enrichment for CTCF, RAD21, SMC3, and ZNF143 as did those trained and predicted on the same cell line (Figure S12-S13). These results indicate that *preciseTAD* pre-trained models can be successfully used to accurately predict domain boundaries for cell lines lacking Hi-C data but for which genome annotation data is available.

### Boundary predictions across cell lines

Given the success of *preciseTAD* in predicting domain boundaries across cell lines, we investigated the possibility of predicting boundaries for cell lines with CTCF/RAD21/SMC3/ZNF143 genome annotation data. The ENCODE project provides information about 132 human cell lines; however, only 4 of them have all four annotations and 48 have none (Figure S14A, Table S6). Therefore, we considered Avocado, a deep learning method providing pre-trained models to impute missing genomic annotations for 400 cell- and tissue types.^41^ Avocado was able to predict the location of three transcription factors (CTCF/RAD21/SMC3) in 60 cell lines available in ENCODE. We found that the signal distributions for all three factors around the midpoints of experimental ENCODE and Avocado-predicted TFBSs are comparable, and the Avocado-predicted CTCF sites are a subset of of ENCODE CTCF sites (Figure 6A). These results suggest that Avocado-predicted genomic annotations can be used when cell type-specific ENCODE data is missing.

**Figure 6.**
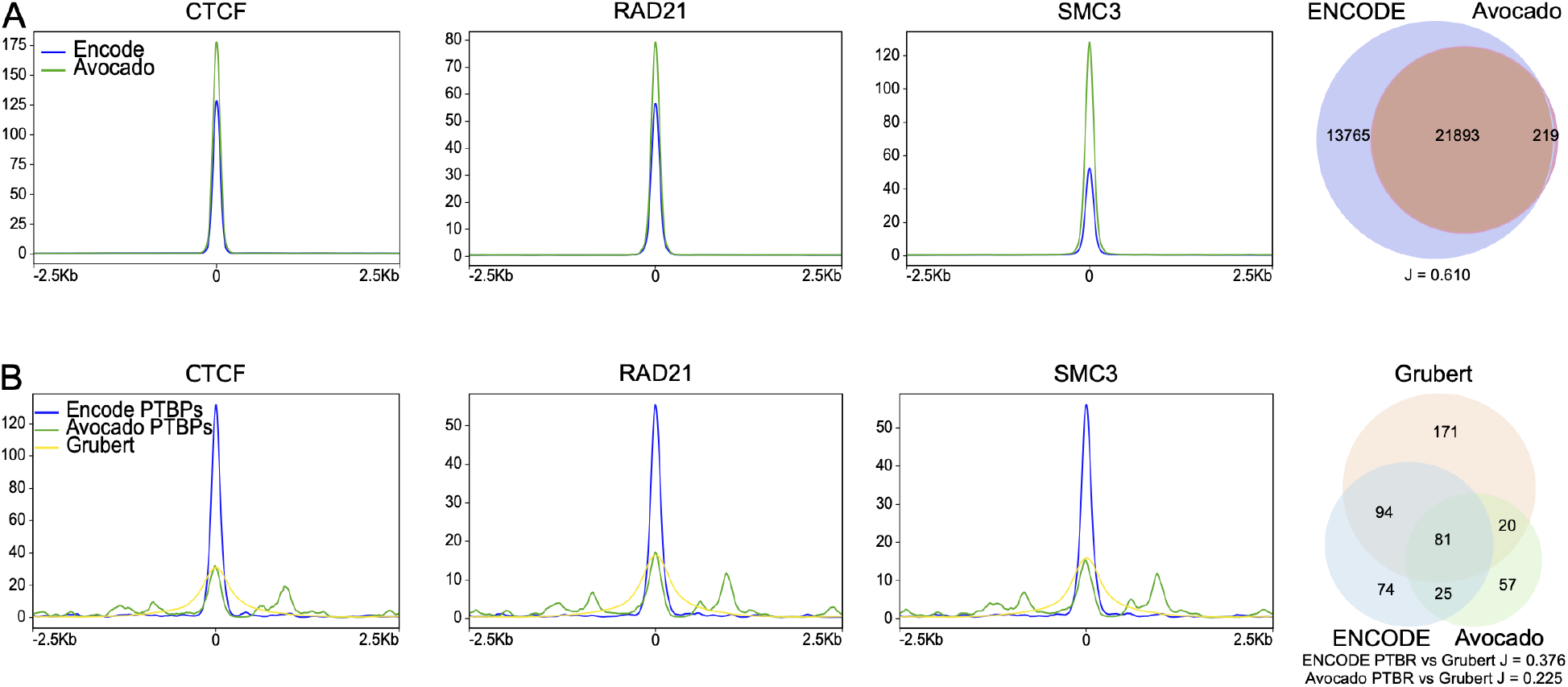
Avocado-imputed genomic annotations can be used for *preciseTAD* predictions. (A) Signal distribution around the midpoint of ENCODE and Avocado TFBSs and the Venn diagram of overlap between ENCODE and Avocado CTCF binding sites. (B) Signal distribution around PTBPs predicted from ENCODE and Avocado data and the midpoints of Grubert anchors, and the Venn diagram of overlaps between ENCODE/Avocado-predicted PTBPs and the Grubert data. Data for GM12878 cell line are shown. For the Venn diagrams, regions were flanked by 5kb.

We retrained *preciseTAD* models to use CTCF/RAD21/SMC3 annotations. Using three instead of four transcription factors resulted in a non-significant drop in performance (Figure S14B), suggesting ZNF143 is less critical for boundary prediction. We applied those models to predict domain boundaries in 60 cell lines, using Avocado-predicted genomic annotations when ENCODE data was unavailable. To examine the predictive power of Avocado genomic annotations, we compared PTBRs and PTBPs predicted from ENCODE and Avocado annotations (Figure 6B). We found that the signal distribution around PTBPs predicted using Avocado data was less than that of ENCODE but remained comparable with that observed from Grubert data (Figure 6B). Similarly, Avocado PTBPs showed less overlap with Grubert data (Figure 6B). The inferior performance of Avocado-predicted genomic annotations is expected as Avocado has been reported to perform less optimally in imputing transcription factors.^41^ In summary, we demonstrate that *preciseTAD* predicts boundaries comparable with those observed experimentally and provides base-level domain boundary predictions for 60 cell lines.

## Discussion

We present *preciseTAD*, a transfer learning approach for the precise prediction of TAD and chromatin loop boundaries from functional genomic annotations. *preciseTAD* leverages a random forest (RF) classification model built on boundaries obtained from low-resolution chromatin conformation capture data, and high-resolution genomic annotations as the predictor space. *preciseTAD* predicts the probability of each base being a boundary, and identifies the precise location of boundary regions and the most likely boundary points. We performed extensive optimization of our RF model by systematically comparing different Hi-C data resolutions, feature engineering procedures, and resampling techniques. Our results demonstrate that distance between boundary regions and genomic annotations coupled with random under-sampling results in the best model performance. We show that binding of four transcription factors (SMC3, RAD21, CTCF, ZNF143) is sufficient for accurate boundary predictions. Compared with ChIA-PET-detected cohesin-mediated loops,^44^ we showed that *preciseTAD*-predicted boundaries better agree with biological properties of experimental data. Models trained in one cell type can accurately predict boundaries in another cell type without Hi-C data, requiring only cell type-specific genomic annotations. *preciseTAD* is implemented as an R package, while pre-trained models for predicting domain boundaries using genomic annotation data are provided via an ExperimentHub R package *preciseTADhub*. We provide a resource of predicted boundaries for 60 cell lines using Avocado-imputed genomic annotation data for cells lacking experimental data.

*preciseTAD* allows for predicting any type of 3D features observed in chromatin conformation capture data. Emerging evidence suggests the existence of different types of 3D domains and domain boundaries.^7,8^ Consequently, subpopulations of 3D domains can be defined by optimizing different sets of computational and biological characteristics (e.g., enrichment of CTCF binding motifs, high occupancy of CTCF/RAD21/H3K36me3 at boundaries, reproducibility, high intra-vs. inter-TAD difference in contact frequencies),^49^ using different training data (e.g., CTCF- and RNAPII ChIA-PET),^35^ or distinguishing long-range cohesin-dependent and short-range cohesin-independent domains.^18,50^ Boundaries have also been defined by the patterns of CTCF orientation,^48^ actively transcribed regions,^51^ and the level of hierarchy.^52–54^ Recent research distinguishes CTCF-associated boundaries, CTCF-negative YY1-enriched boundaries, CTCF- and YY1-depleted promoter boundaries, and the fourth class of weak boundaries largely depleted of all three features.^55^ *preciseTAD* can be trained on boundaries defined by other algorithms and characteristics. Our future work will include incorporating the directionality of CTCF binding in predictive modeling, including additional predictor types, and defining separate models trained on different boundary types defined by different technologies.

In summary, we demonstrate that domain boundary prediction is a multi-faceted problem requiring consideration of multiple statistical and biological properties of genomic data. Simply considering properties of Hi-C contact matrices ignores the fundamental roles of known molecular drivers of 3D chromatin structures. Instead, we propose *preciseTAD*, a supervised machine learning framework that leverages both Hi-C contact matrix information and genomic annotations. Our method introduces three concepts - *shifted binning*, *distance*-type predictors, and *random undersampling* - which we use to build random forest classification models for predicting boundary regions. Our method can bridge the resolution gap between 1D genomic annotations and 3D Hi-C data for more precise and biologically meaningful boundary identification. We introduce *preciseTAD*, an open source R package for leveraging random forests to predict domain boundaries at base-level resolution, as well as the genomic coordinates of predicted boundaries for 60 cell types. We hope that *preciseTAD* will serve as an efficient and easy-to-use tool to further explore the genome’s 3D organization.

## Methods

### Data sources

TAD and loop boundaries called by *Arrowhead*^56^ and *Peakachu*^17^ tools were used as training and testing data. The autosomal genomic coordinates in the GRCh37/hg19 human genome assembly were considered. Arrowhead-defined TAD boundaries were called from Hi-C data for the GM12878 and K562 cell lines (MAPQ>0, 5 kb, 10 kb, 25 kb, 50 kb, and 100 kb resolutions) using the Arrowhead tool from Juicer^56^ with default parameters. Peakachu chromatin loop boundaries for GM12878 and K562 cell lines were downloaded from the Yue lab website. Experimentally obtained (ChIA-PET) cohesin-mediated chromatin loops, which we refer to as *Grubert* data, were obtained from the Supplementary Table 4 of^44^ (Table S2). Unique boundaries were considered as the midpoints within the coordinates of each chromatin loop anchor. Chromosome 9 was excluded due to the inability to call Arrowhead domains at 5 kb and 10 kb resolutions for the K562 cell line (high sparsity), unless specified otherwise. Cell-line-specific genomic annotations (BroadHMM chromatin states (BroadHMM), histone modifications (HM), and transcription factor binding sites (TFBS)) were obtained from the UCSC Genome Browser Database(Table S2).

### Shifted-binning for binary classification

In Hi-C, each chromosome is binned into non-overlapping regions of length *r*, typically, 5 kb and above. The *r* parameter defines the resolution of Hi-C data. Here, we designed a strategy called *shifted binning* that partitions the genome into regions of the same length *r*, but with middle points corresponding to boundaries defined by the original binning. To create shifted binning, the first shifted bin was set to start at half of the resolution *r* and continued in intervals of length *r* until the end of the chromosome (*mod r* + *r/2*). The shifted bins, referred hereafter as bins for simplicity, were then defined as boundary-containing regions (*Y = 1*) if they contained a TAD (or loop) boundary, and non-boundary regions (*Y = 0*) otherwise, thus establishing the binary response vector (**Y**) used for classification (Figure S2A).

### Feature engineering

Cell line-specific genomic annotations were used to build the predictor space. Bins were annotated by one of either the average signal strength of the corresponding annotation (*Peak Signal Strength*), the number of overlaps with an annotation (*Overlap Count (OC)*), the percent of overlap between the bin and the total width of genomic annotation regions overlapping it (*Overlap Percent (OP)*), or the distance in bases from the center of the bin to the center of the nearest genomic annotation region (*Distance*) (Figure S2B). A (*log*_2_ + 1)-transformation of distance was used to account for the skewness of the distance distributions (Figure S3). Models built using a *Peak Signal Strength* predictor space were only composed of histone modifications and transcription factor binding sites as BroadHMM chromatin states lack signal values.

### Addressing class imbalance

To assess the impact of class imbalance (CI), defined as the proportion of boundary regions to non-boundary regions, we evaluated three resampling techniques: *Random Over-Sampling (ROS)*, *Random Under-Sampling (RUS)*, and *Synthetic Minority Over-Sampling Technique (SMOTE)*. For ROS, the minority class was sampled with replacement to obtain the same number of data points in the majority class. For RUS, the majority class was sampled without replacement to obtain the same number of data points in the minority class. For SMOTE, under-sampling was performed without replacement from the majority class, while over-sampling was performed by creating new synthetic observations using the *k* = 5 minority class nearest neighbors^46^ (implemented in the `DMwR` v.0.4.1 R package). We restricted the SMOTE algorithm to 100% over-sampling and 200% under-sampling to create perfectly balanced classes.

### Establishing optimal data level characteristics for boundary region prediction

Random forest (RF) classification models (the caret v.6.0 R package)^57^ were compared between combinations of data resolutions, feature engineering procedures, and resampling techniques. Following recommendations to evaluate the model on unseen data,^58^ a *holdout chromosome* technique was used for estimating model performance. The *i^th^* holdout chromosome was identified and a data matrix, *A*_*N*×(*p*+1)_, was constructed by combining the binned genome from the remaining chromosomes (1, 2,…, *i* – 1, *i* + 1,…, 21, 22), where *N* = [*n*_1_*n*_2_… *n*_21_*n*_22_]′ and *n_k_* is the length of chromosome *k* after being binned into non-overlapping regions of resolution *r*, such that *k* ≠ *i*. The number of annotations, *p*, and the response vector, **Y**, defined the column-wise dimension of the matrix *A*. Re-sampling was then performed on *A*, and a RF classifier was trained using 3-fold cross-validation to tune for the number of annotations to consider at each node (*mtry*). The number of trees (*ntree*) that were aggregated for each RF model was set to 500. The minimum number of observations per root node (*nodesize*) was set to 0.1% of the rows in the data. The binned data for the holdout chromosome *i* was reserved for testing. The response vector associated with the testing data (**Y***_test_*) was built using Grubert-defined chromatin loops as a ground truth when validating the models. Models were then evaluated using Balanced Accuracy (BA), defined as the average of sensitivity and specificity:

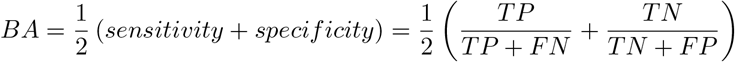

where values of the confusion matrix, true negatives (TP), false positives (FP), true negatives (TN), and false negatives (FN) were related to genomic bins that contained a Grubert-defined boundary in the test data. That is, TP refers to the number of bins correctly identified as containing a boundary (true positives), FP refers to the number of bins incorrectly identified as containing a boundary (false positives), TN refers to the number of bins correctly identified as not containing a boundary (true negatives), and FN refers to the number of bins incorrectly identified as not containing a boundary (false negatives). Each of these quantities is obtained from the confusion matrix created by validating the model on the test data. The process was repeated for each *i^th^* holdout chromosome, and performances were aggregated using the mean and standard deviation.

### Feature selection and predictive importance

Many genomic annotations, notably architectural proteins, tend to exhibit an extensive pattern of colocalization (correlation). To suitably reduce the predictor space and improve computational efficiency, while maintaining optimal performance, we utilized recursive feature elimination (RFE). We estimated the near-optimal number of necessary features, ranging from 2 to the maximum number of features incremented by the power of 2. We then aggregated the predictive importance of the union of the optimal set of features across holdout chromosomes using the mean decrease in node impurity among permuted features in out-of-bag samples to determine the most common and top-ranked annotations for predicting boundary regions.

### Evaluating performance across cell lines

We used the same holdout chromosome strategy to evaluate a model trained in one cell line on unseen data from another cell line.^58^ Given two cell lines, GM12878 and K562, we first evaluated the performance of cell line-specific models. That is, models trained on cell line-specific data from *n* – 1 chromosomes were evaluated on the *i^th^* holdout chromosome data from the same cell line. Second, we evaluated models trained on cell line-specific data from *n* – 1 chromosomes using the *i^th^* holdout chromosome data from a different cell line. That is, models trained using K562 cell line-specific data were evaluated on unseen chromosome data from the GM12878 cell line. This process was repeated for each holdout chromosome. To evaluate performance, we constructed receiver operating characteristic (ROC) curves composed of the average sensitivities and specificities at different cutoffs, across each holdout chromosome, and reported the corresponding average area under the curve (AUC). As before, the response vector for the test data was derived from cell line-specific Grubert boundaries.

### Boundary prediction at the base-level resolution using *preciseTAD*

To investigate whether we could alleviate the resolution limitations of conventional domain calling tools, we developed *preciseTAD*. This algorithm leverages an optimized random forest model in conjunction with density-based and partitioning techniques to predict boundaries at base-level resolution (Figure S7). A random forest model (*M*) was built on the optimal combination of predictor type (predictor space), resampling technique (class balancing correction), and top-ranked annotations (most informative genomic elements) for a set of binned chromosomes {*k*|*k* ≠ *i*}. To precisely identify boundary locations, we first constructed a *base-level resolution* predictor space for the chromosome *i*, *A*_*n*×*p*_, where *n* is the length of chromosome *i* and *p* is the optimal number of annotations. We then evaluate *M* on the base-level predictor space to extract the probability vector, *π_n_*, denoting each base’s probability of being a boundary. A threshold *t* specifies the probability at which a base with *π_n_* ≥ *t* is designated as a potential boundary (the default *t* =1). Next, density-based spatial clustering of applications with noise (DBSCAN; version 1.1-5) is applied to the matrix of pairwise genomic distances between boundary-annotated bases, *D*, such that *π_n_* ≥ *t*. The minimum and maximum coordinates of each cluster, *k*, of spatially colocalized bases were termed *preciseTAD boundary regions* (PTBR). To precisely identify a single base among each PTBR, *preciseTAD* implements partitioning around medoids (PAM) on the distance matrix, *D_k_*, derived from each cluster. The corresponding cluster medoid was defined as a *preciseTAD boundary point* (PTBP), making it the most representative base coordinate within each clustered PTBR.

The DBSCAN algorithm has two parameters, *MinPts* and *eps* (*ϵ*). The *MinPts* parameter was set to the recommended value of 3, representing 1 + *dim*(*data*).^59^ To decide on the optimal value of *t* and *ϵ* in *preciseTAD*, we considered the normalized enrichment (*NE*) of flanked boundaries. *NE* was calculated as the average number of overlaps between genomic annotations (CTCF, RAD21, SMC3, and ZNF143) and flanked boundaries, divided by the total number of boundaries. The rationale here is to find a combination of parameters producing the largest number of overlaps between predicted boundaries and genomic annotations. We evaluated *NE* for combinations of *t* = {0.975, 0.99, 1.0} and *ϵ* = {1000, 5000, 10000, 15000, 20000, 25000}. The heuristic of *ϵ* is that density-reachable bases with genomic distances less than *ϵ* should occupy the same designated cluster. The default combination was set to *t* = 1.0 and *ϵ* = 10000 based on our tests (Figure S15).

### Evaluating called and predicted boundary precision

We assessed the biological significance of our predicted boundaries by their association with the signal of CTCF, RAD21, SMC3, and ZNF143 using *deepTools* (version 2.0)^60^ (*computeMatrix*, *plotProfile* tools). Additionally, we compared the median *log*_2_ genomic distances between TAD boundaries and the same top predictive ChIP-seq annotations using Wilcoxon Rank-Sum tests. Furthermore, we compared the overlap between predicted and called boundaries in GM12878 and K562 cell lines. Boundaries were first flanked by resolution, *r*, and overlaps were visualized using Venn diagrams from the `Vennerable` R package (version 3.1.0). Overlaps were further quantified using the Jaccard index defined as

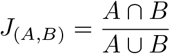

where A and B represent genomic regions created by flanked boundaries. All statistical analyses were performed in R (version 4.0.1). The significance level was set to 0.05 for all statistical tests.

### Predicting boundaries across cell lines

We implemented a strategy to predict domain boundaries across cell lines. To do so, we trained *preciseTAD* models on boundaries and genomic annotation data from one cell line and used it to predict boundaries using genomic annotation data from another cell line. Results were compared by assessing the overlap between flanked same-cell-line and cross-cell-line predicted boundaries using Venn diagrams, Jaccard indices, and signal distribution plots.

To predict boundaries in 60 cell lines, we first compiled a set of cell line-specific genomic annotations (CTCF, RAD21, SMC3). For cells lacking some or all genomic annotations, we used Avocado v.0.3.6^41^ to impute missing annotations. Avocado imputes signal profiles as bigWig files which we converted to bedGraph and called peaks using UCSC’s bigwigtobedgraph and MACS2 v.2.2.7.1 with default settings. We note that Avocado can impute data for only three transcription factors, CTCF, RAD21, SMC3. Therefore, we retrained *preciseTAD* models using these three transcription factors in GM12878 cell line and Arrowhead/Peakachu boundaries. For Avocado predictions, the following cell line names were manually matched: H1 = H1-hESC, H7 = H7-hESC. Predictions were made using the hg38 genome assembly; therefore, all data were either downloaded as hg38 or lifted over to hg38 genome assembly.

## Data Availability

All datasets used in this work are summarized in Tables S1, S2, S6. The predicted domain boundary regions and points for 60 cell lines can be downloaded from https://dozmorovlab.github.io/preciseTAD.

## Code availability

*preciseTAD* is available on Bioconductor https://bioconductor.org/packages/preciseTAD/ and GitHub under the MIT license https://github.com/dozmorovlab/preciseTAD/

## Author contributions

MD conceptualized and supervised the study, and wrote the manuscript with input from all authors. SS and MD wrote the software associated with the R package. SS performed the analyses. MM performed additional analyses and generated figures.

## Acknowledgements

We would like to thank Drs. Jonathan Wren and Cory Giles for helpful comments.

