## Supplementary material for "preciseTAD: A transfer learning framework for 3D domain boundary prediction at base-pair resolution": Figure S1

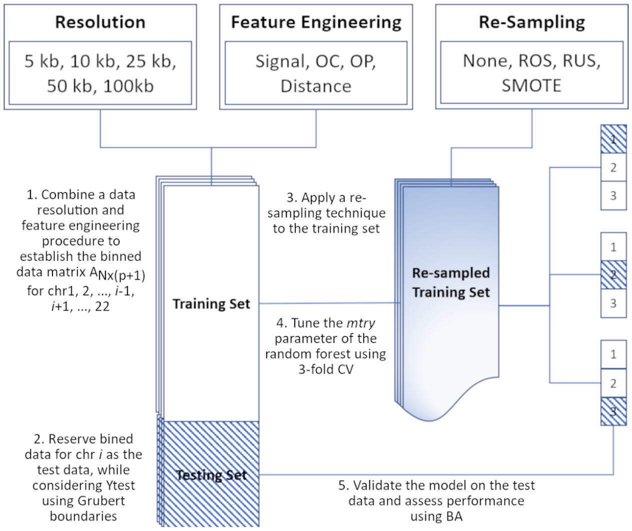


**Figure S1. A machine learning framework for optimizing domain boundary region prediction models.** Step 1 combines the response vector ($\mathbf{Y}_N$) from *shifted binning* and a feature engineering procedure to form the data matrix $A_{N\times(p+1)}$ for chromosomes $\{1,2,...,i-1,i+1,...,22\}$. Step 2 reserves the predictor-response matrix for the holdout chromosome $i$ as the test data using Grubert-defined boundaries as $\mathbf{Y}_{test}$. Step 3 applies a resampling technique to the training data to address the class imbalance. Step 4 trains the random forest model and performs 3-fold cross-validation to tune the mtry parameter. Finally, step 5 validates the model on the separate test data composed of the binned data from the holdout chromosome $i$ and evaluates the model performance using balanced accuracy (BA, see Methods).

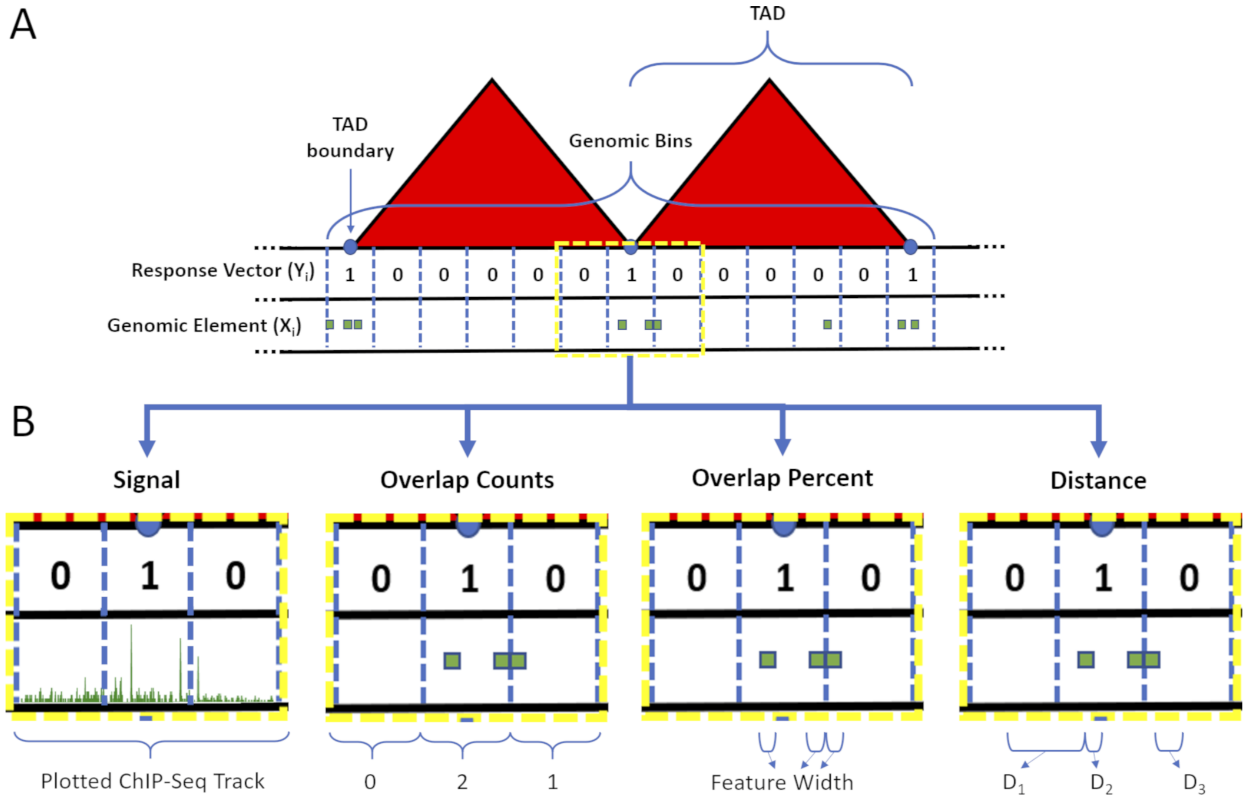


**Figure S2. Resolution-specific data construction and feature engineering for random forest modeling.** (A) The linear genome was binned into non-overlapping resolution-specific intervals using *shifted binning* (see Methods). The response vector **Y** was defined as 1/0 if a genomic bin overlapped/did not overlap with a TAD (or loop) boundary. (B) Four types of associations between bins (blue dashed lines) and genomic annotations (green shapes) were considered to build the predictor space, including Average Peak Signal (Signal), Overlap Counts (OC), Overlap Percent (OP), and $log_2$ distance (Distance).

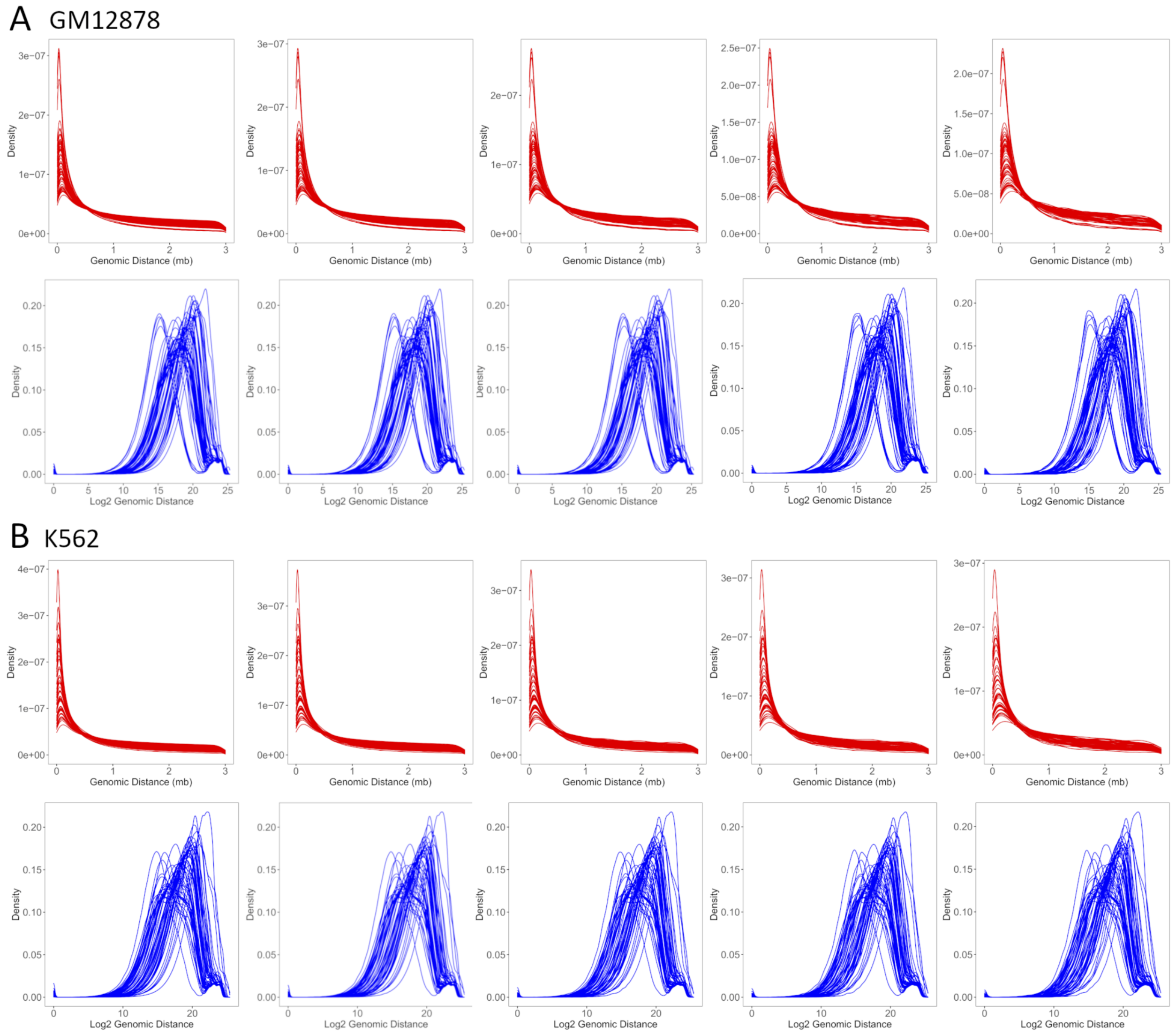


**Figure S3. The $log_2$ transformation of genomic distances normalizes their distributions.** Distances are measured as the number of bases from the center of a genomic bin to the nearest genomic annotation center. Density curves of distances before (red) and after (blue) performing a $log_2$ transformation across 5 kb, 10 kb, 25 kb, 50 kb, and 100 kb data resolutions for both the (A) GM12878 and (B) K562 cell lines. Each density curve represents an individual genomic annotation (77 total).

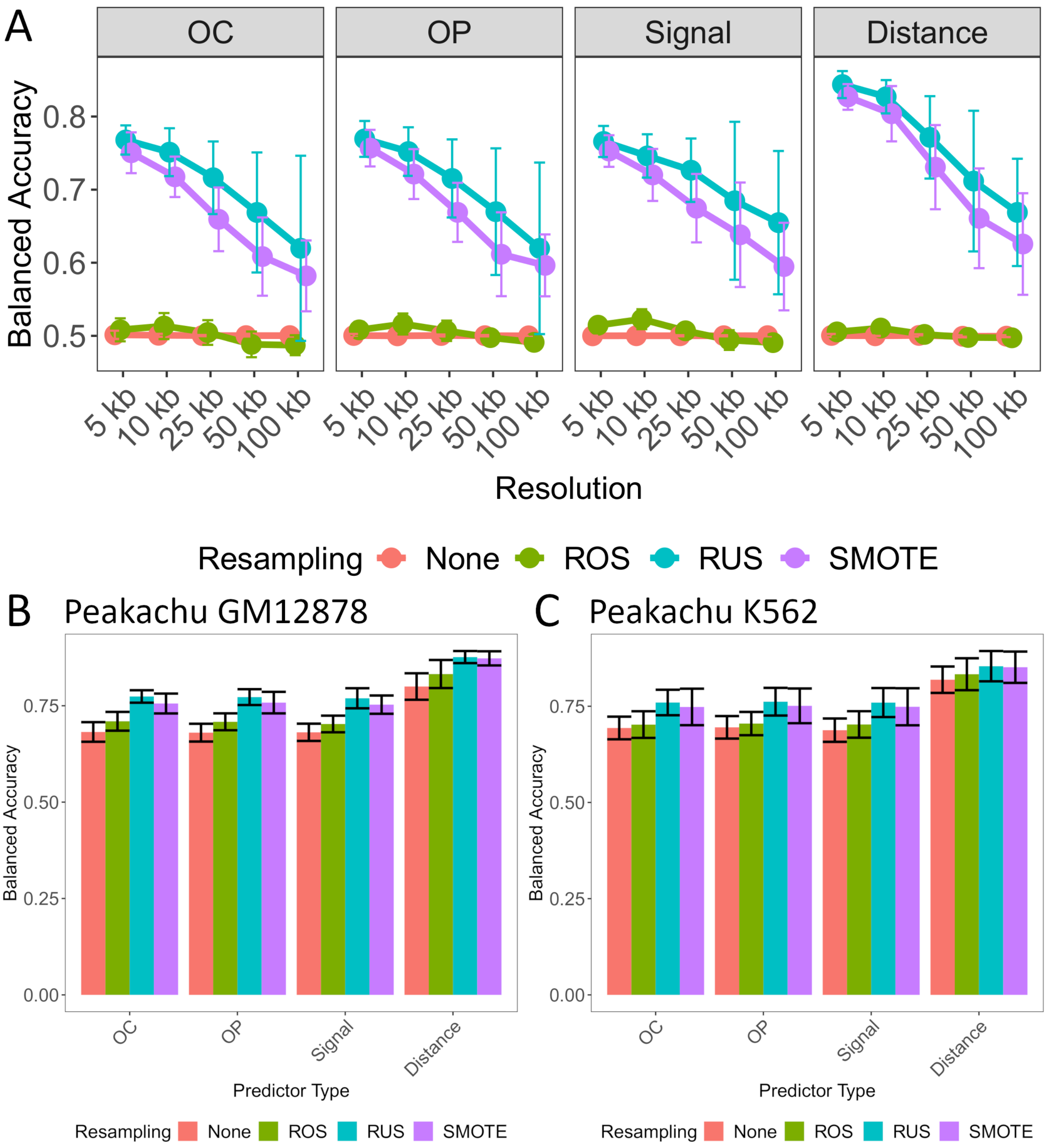


**Figure S4. Determining optimal data level characteristics for building TAD boundary region prediction models on K562.** (A) Averaged balanced accuracies are compared across resolution, within each predictor-type: overlap count (OC), overlap percent (OP), average Signal and Distance and across resampling techniques: no resampling (None; red), random over-sampling (ROS; green), random under-sampling (RUS; blue), and synthetic minority over-sampling (SMOTE; purple) when using Arrowhead ground truth boundaries for K562. Averaged balanced accuracies are compared for Peakachu-trained models built on (B) GM12878 and (C) K562 within each predictor-type: OC, OP, Signal and Distance, and across resampling technique: no resampling (None; red), random over-sampling (ROS; green), random under-sampling (RUS; blue), and synthetic minority over-sampling (SMOTE; purple). Error bars indicate 1 standard deviation from the mean performance across each holdout chromosome used for testing.

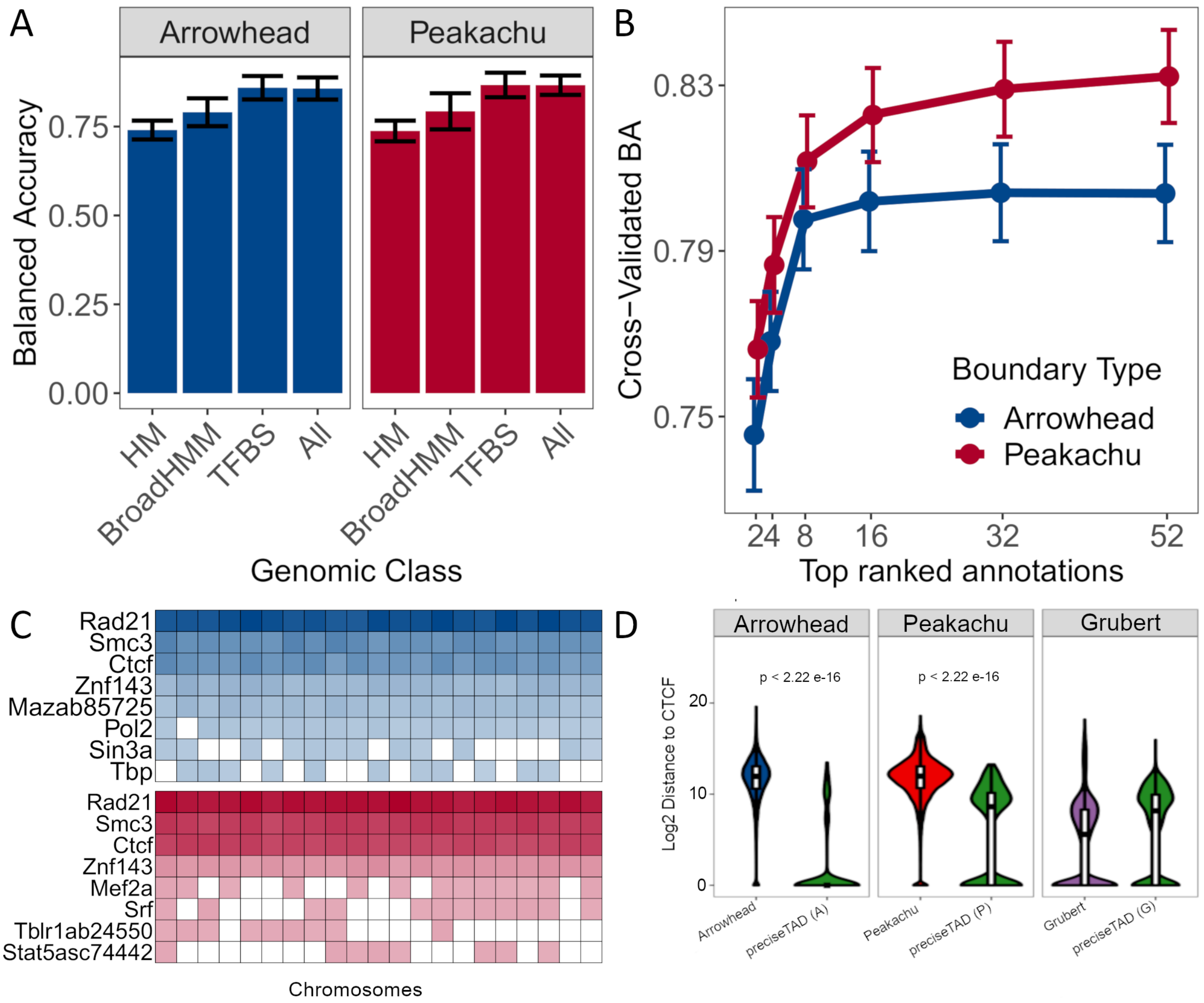


**Figure S5. SMC3, RAD21, CTCF, and ZNF143 transcription factors accurately predict TAD and loop boundaries in K562.** (A) Barplots comparing performances of TAD (Arrowhead) and loop (Peakachu) boundary prediction models using histone modifications (HM), chromatin states (BroadHMM), transcription factor binding sites (TFBS), in addition to a model containing all three classes (ALL). (B) Recursive feature elimination (RFE) analysis used to select the optimal number of predictors. Error bars represent 1 standard deviation from the mean cross-validated accuracy across each holdout chromosome. (C) Clustered heatmap of the predictive importance for the union of the top 8 most predictive chromosome-specific TFBSs. The columns represent the holdout chromosome excluded from the training data. Rows are sorted in decreasing order according to the columnwise average importance. (D) Violin plots illustrating the $log_2$ genomic distance distribution from original Arrowhead/Peakachu boundaries vs. *preciseTAD*-predicted boundaries to the nearest CTCF sites. The p-values are from the Wilcoxon Rank Sum test.

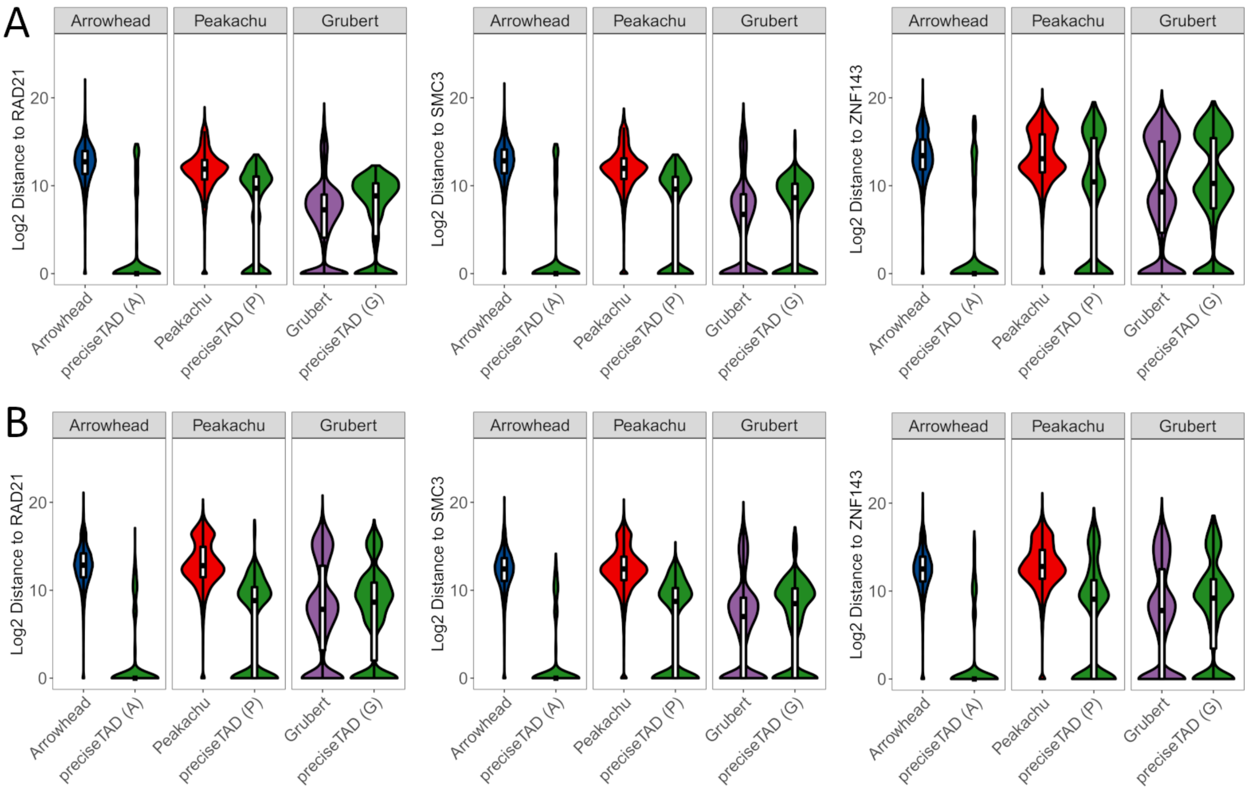


**Figure S6. *preciseTAD* boundaries are spatially closer to known molecular drivers of 3D chromatin.** Violin plots illustrating the $log_2$ genomic distance distribution from original Arrowhead/Peakachu boundaries vs. *preciseTAD*-predicted boundaries to the nearest RAD21/SMC3/ZNF143 sites. Data for (A) GM12878 and (B) K562 cell lines are shown.

1. Consider an optimized RF model (M) built on the set of autosomal chromosomes $\{k|i \notin k\}$ binned at some resolution $r$

2. **for** *each chr i* **do**

    3. Construct the base-level resolution predictor space $A_{n\times p}$ where $n$ is the length of chr $i$ and $p$ is the number of predictors

    4. Assign threshold $\{t|0 \leq t \leq 1\}$ and $\{\epsilon|\epsilon > 0\}$

    5. **if** $|t| > 1$ *or* $|\epsilon| > 1$ **then**

        6. **for** *each combination (l) of t and* $\epsilon$ **do**

            7. Evaluate M on $A_{n\times p}$ to get the probability of each genomic coordinate as being a domain boundary $\pi_n$

            8. Subset $\{\pi_n|\pi_n \geq t_l\}$

            9. Construct the pairwise distance matrix $D$ between genomic coordinates where $\pi_n \geq t_l$

            10. Apply DBSCAN on $D$ with $MinPts = 3$ and $eps = \epsilon_l$

            11. **for** *each cluster k identified by DBSCAN* **do**

                12. Assign $w_k$ as the number of coordinates that span each cluster of bases in $k$ (PTBR)

                13. Perform PAM on the sub-distance matrix $D_k$ to extract the cluster medoid $b_k$ (PTBP)

                14. **for** *each predictor p* **do**

                    15. Calculate the normalized enrichment (NE) over all predictors

$$NE = \frac{1}{p}\left[\Sigma_{s=1}^{p}\left[\frac{1}{b}\Sigma_{k=1}^{b} e_{ks}\right]\right]$$

                    where $e_{ks} = \mathbf{I}\{r_s \in (b_k - f, b_k + f)\}$ is the number of elemental regions $r$ of predictor $p$ that overlap with each flanked boundary

                    16. Determine where $NE$ converges as optimal $\{t, \epsilon\}$ combination

                **end**

            **end**

        **end**

        17. Repeat steps 7-14 on $A_{nxp}$ with optimal $\{t, \epsilon\}$

    **else**

        18. Perform steps 7-14 on $A_{nxp}$ such that $t = t_0$ and $eps = \epsilon_0$

    **end**

**end**

**Algorithm 1:** Psuedocode for $preciseTAD$ implementation.

**Figure S7. Pseudocode of the *preciseTAD* algorithm.**

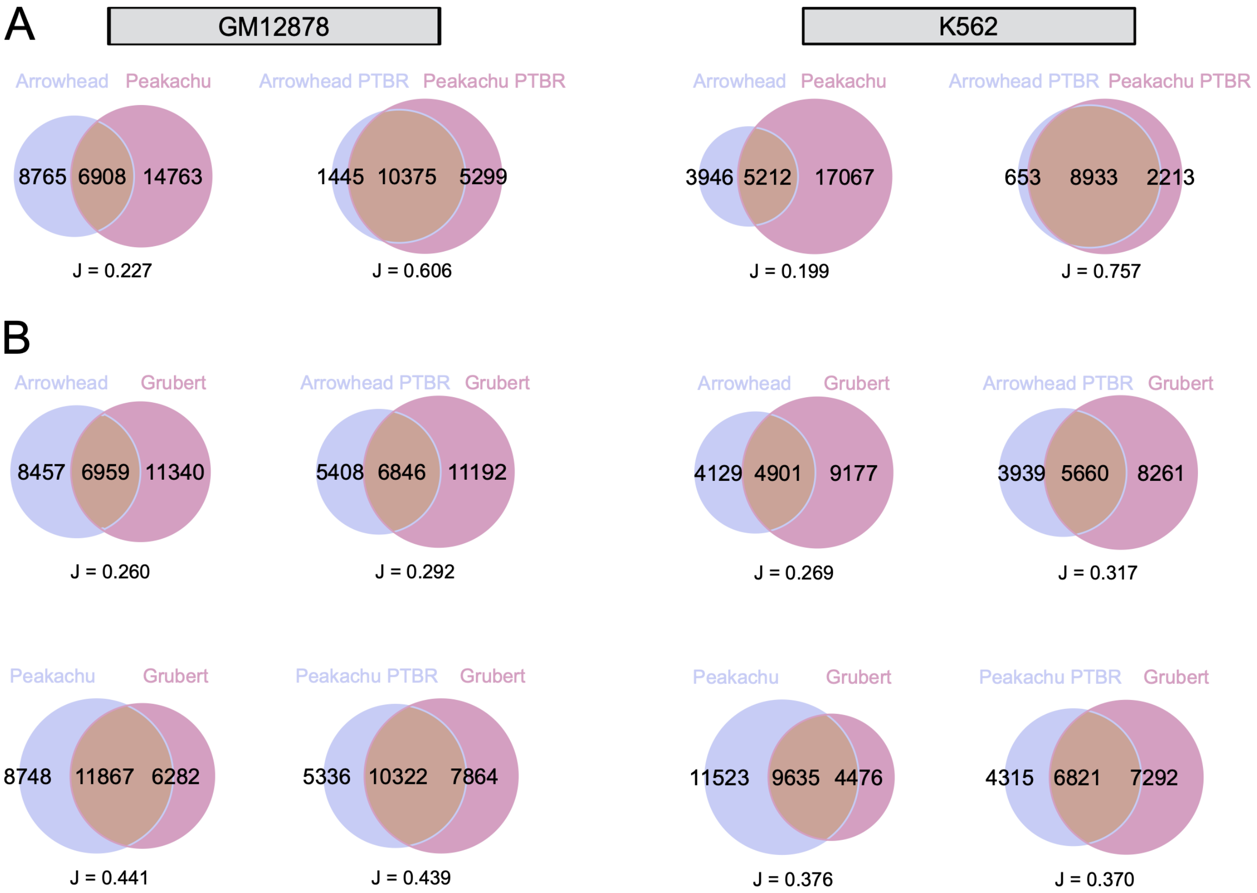


**Figure S8. *preciseTAD* PTBRs show high overlap and agreement with experimental loop boundaries.** Venn diagrams of boundary overlap between (A) original Arrowhead-Peakachu boundaries and Arrowhead-Peakachu PTBRs, and (B) overlaps with Grubert data. Boundaries were flanked by 5 kb.

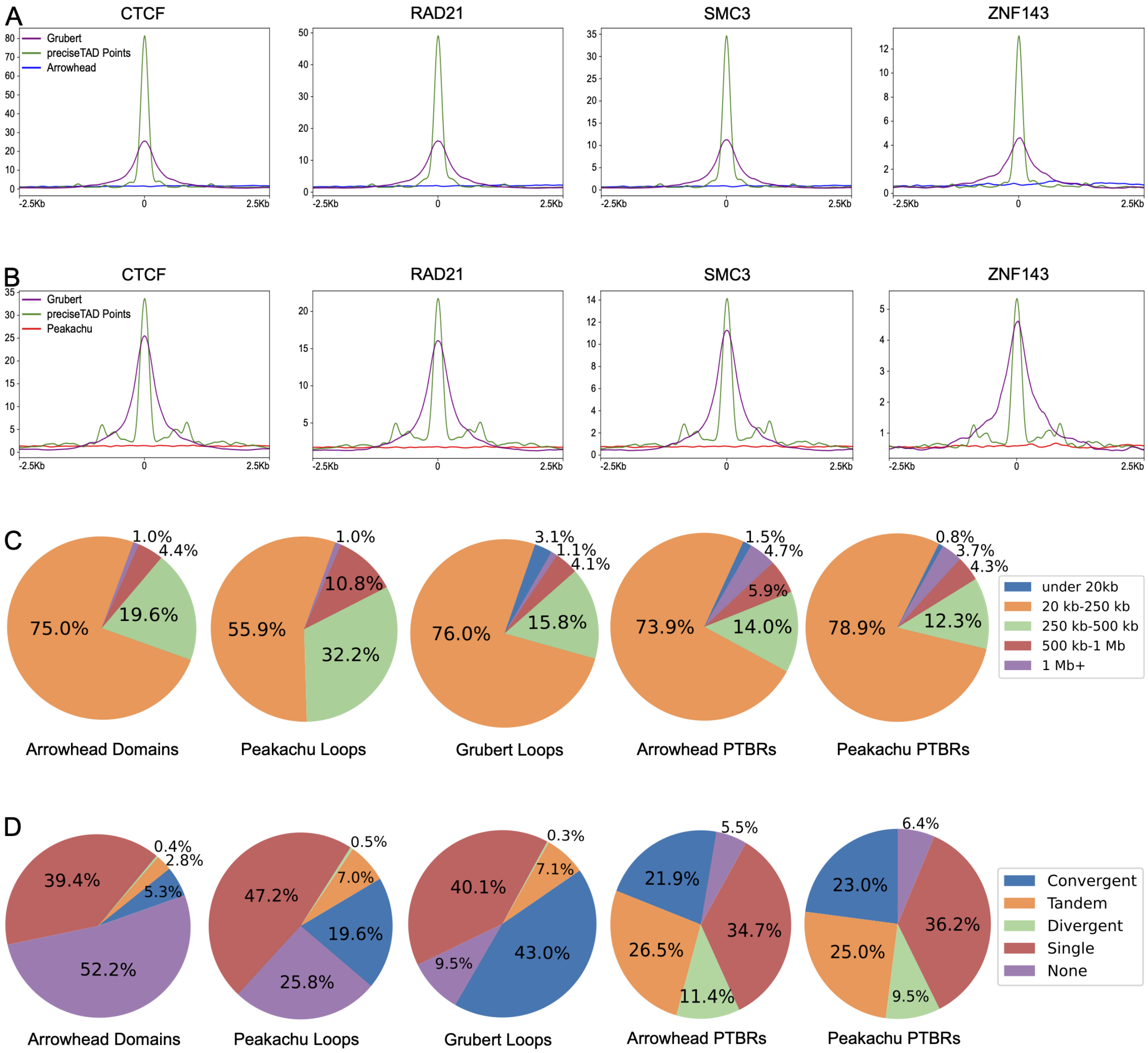


**Figure S9.** ***preciseTAD* boundaries are more enriched for known molecular drivers of 3D chromatin.** Signal enrichment strength of CTCF, RAD21, SMC3, and ZNF143 sites around midpoints of *preciseTAD*-predicted boundaries (green) compared to midpoints of (A) Arrowhead-called boundaries (blue), (B) Peakachu loop boundaries (red). Data for midpoints of Grubert cohesin loop boundaries is shown as a proxy for experimental "ground truth" (purple). Panel inserts show signal enrichment around preciseTAD boundary points vs. Grubert ground truth. (C) Domain size distribution, and (D) CTCF orientation analysis. Data for K562 cell line is shown.

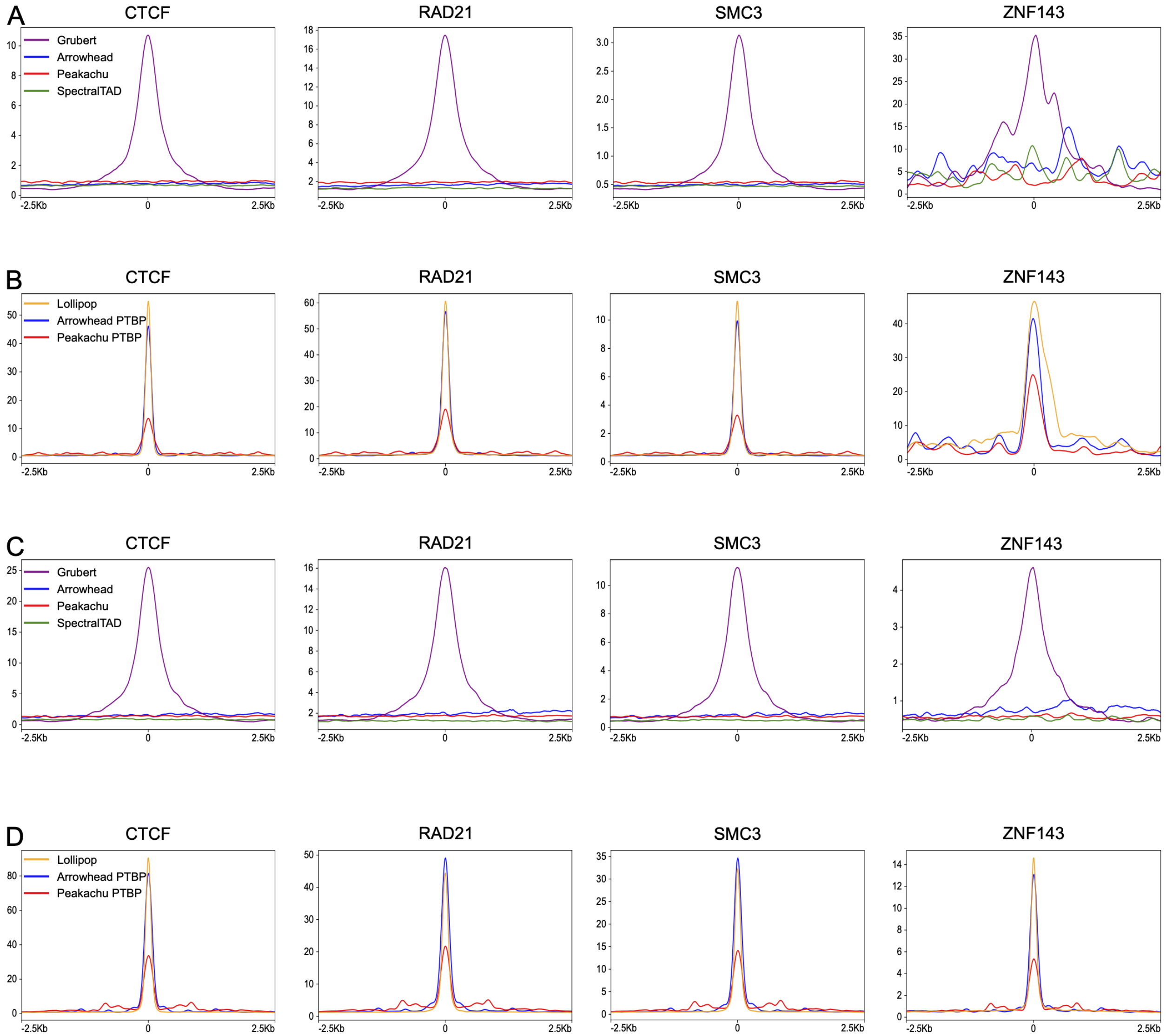


**Figure S10. Comparing enrichment levels between TAD/chromatin loop calling tools.** Signal profile plots comparing the binding strength of top TFBS around Arrowhead (blue), Peakachu (red), SpectralTAD (green) called boundaries vs. experimental Grubert chromatin loop boundaries (purple) in (A) GM12878 and (B) K562 cell lines.

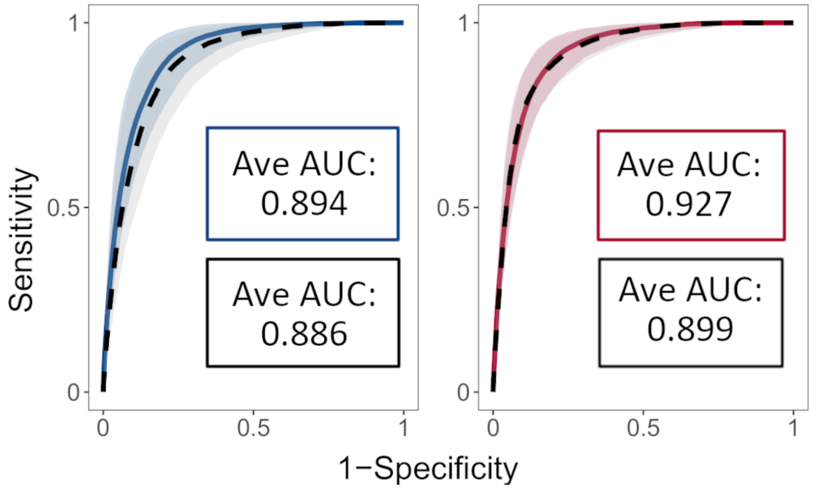


**Figure S11.** ***preciseTAD* models trained in one cell line can accurately predict boundaries in another cell line.** Receiver operating characteristic (ROC) curves and the corresponding average area under the curves (AUCs) when training and testing on K562 data (blue, Arrowhead ground truth; red, Peakachu ground truth) versus training on GM12878 and testing on K5628 data (black, dashed). The curves represent the average sensitivities and specificities across each holdout chromosome. The shaded areas around each curve represent 1 standard deviation from the average.

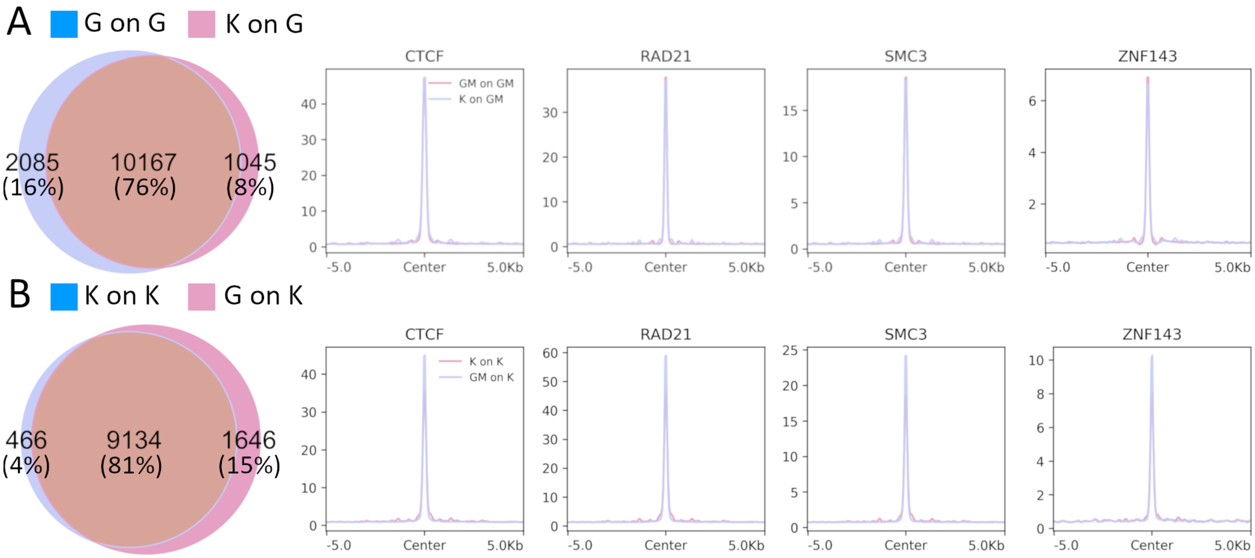


**Figure S12.** ***preciseTAD* trained on Arrowhead accurately predicts boundaries on cell lines using annotation data only.** Venn diagrams and signal profile plots comparing flanked predicted boundaries using Arrowhead trained models. (A) Models trained on GM12878 and predicted on GM12878 (red, GM on GM) vs. models trained on K562 and predicted on GM12878 (blue, K on GM). (B) Models trained on K562 and predicted on K562 (red, K on K) vs. models trained on GM12878 and predicted on K562 (blue, GM on K). Boundaries were flanked by 5 kb.

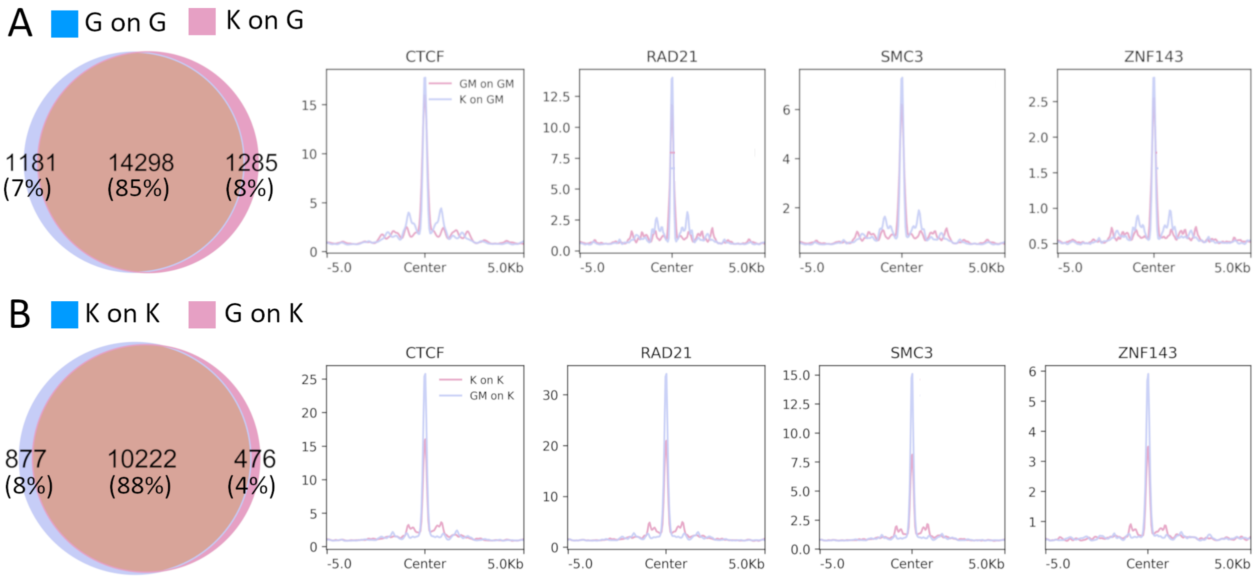


**Figure S13.** ***preciseTAD* trained on Peakachu accurately predicts boundaries on cell lines using annotation data only.** Venn diagrams and signal profile plots comparing flanked predicted boundaries using Arrowhead trained models. (A) Models trained on GM12878 and predicted on GM12878 (red, GM on GM) vs. models trained on K562 and predicted on GM12878 (blue, K on GM). (B) Models trained on K562 and predicted on K562 (red, K on K) vs. models trained on GM12878 and predicted on K562 (blue, GM on K). Boundaries were flanked by 10 kb.

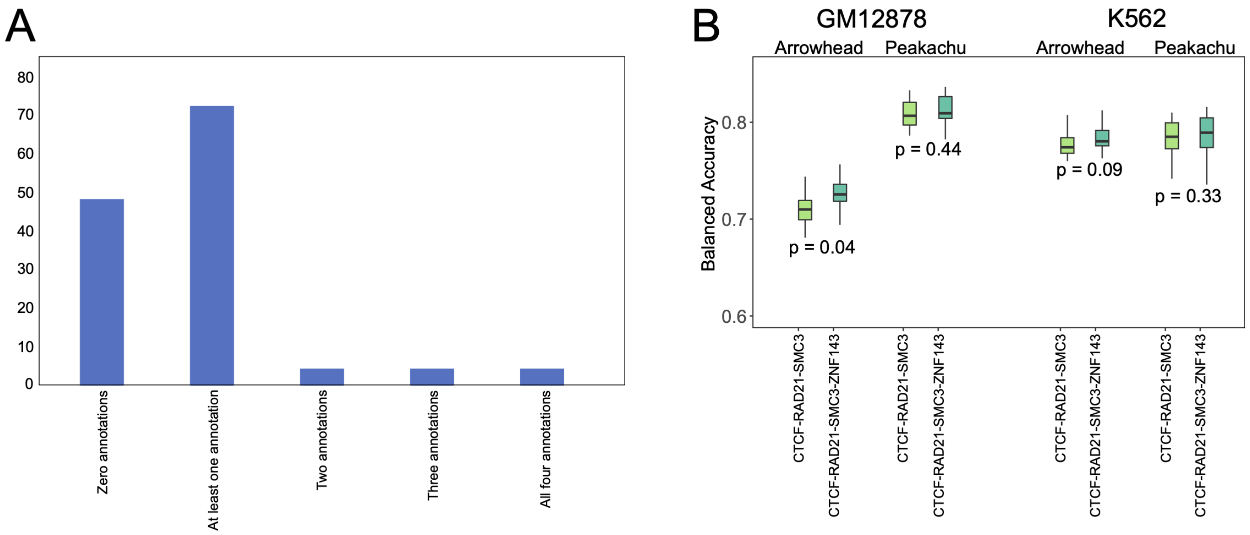


**Figure S14.** (A) Scarcity of cell lines with CTCF/RAD21/SMC3/ZNF143 genomic annotations. (B) Comparable performance of preciseTAD models trained on three vs. four genomic annotations. The p-values are from the Wilcoxon Rank Sum test.

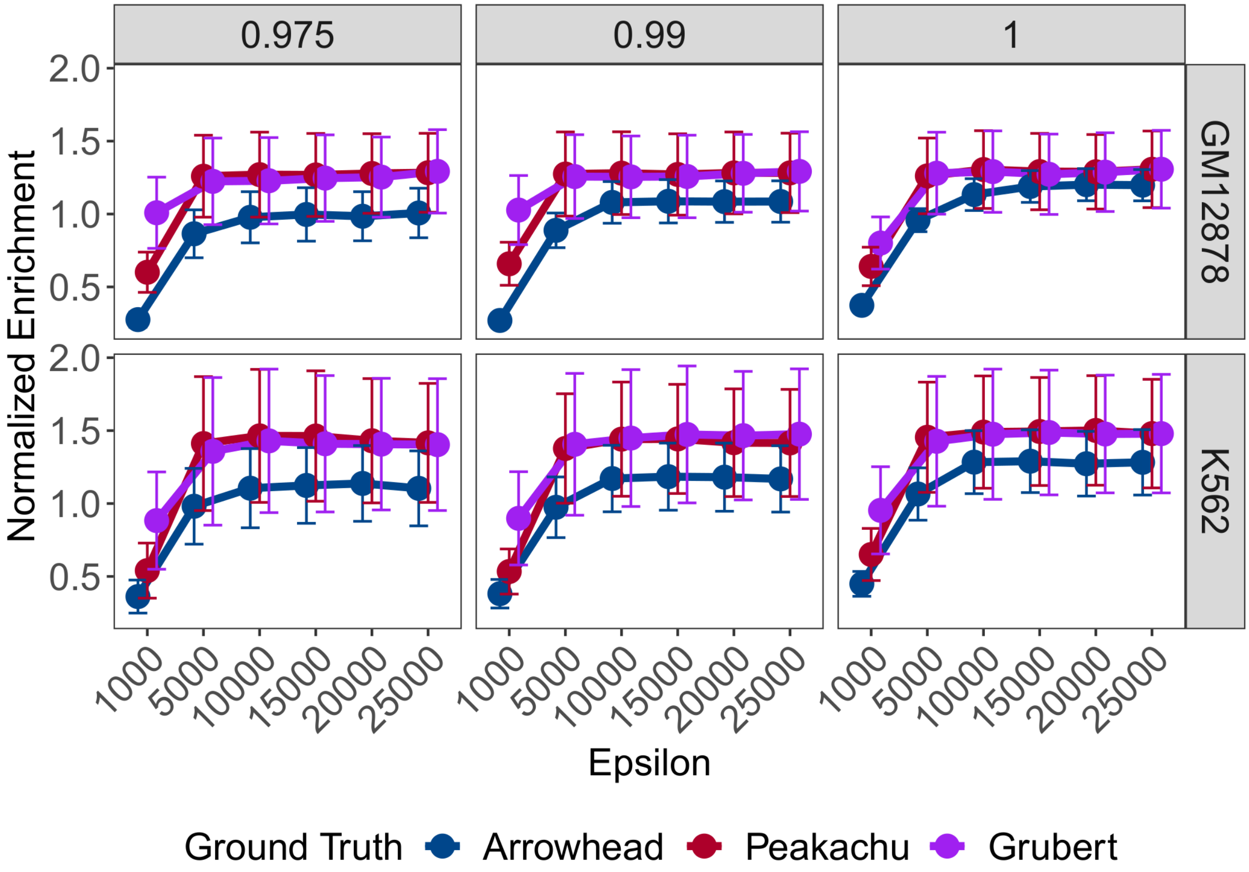


**Figure S15. Normalized Enrichment levels suggest t=1.0 and $\epsilon$=10000 as the most optimal parameters for biologically relevant *preciseTAD*-predicted boundaries.** Linecharts illustrating the normalized enrichment (NE, see Methods) among resolution-flanked *preciseTAD*-predicted boundaries for different combinations of thresholds ($t$) and epsilon-neighborhood parameter values ($\epsilon$). Error bars indicate 1 standard deviation from the mean.

**Table S1 .** Data sources for Hi-C matrices used to call topologically associating domains with Arrowhead, as well as chromatin loop boundaries obtained by Peakachu and Grubert.

| Publisher | Tool | Library | Cell.line | Source |
|---|---|---|---|---|
| Rao et al | Arrowhead | HIC001-HIC018 | GM12878 | https://www.ncbi.nlm.nih.gov/geo/download/?acc=GSE63525&format=file&file=GSE63525%5FGM12878%5Finsitu%5Fprimary%2Ehic |
| Rao et al | Arrowhead | HIC069-HIC074 | K562 | https://www.ncbi.nlm.nih.gov/geo/download/?acc=GSE63525&format=file&file=GSE63525%5FK562%5Fcombined%2Ehic |
| Salameh et al | Peakachu | | GM12878 | http://promoter.bx.psu.edu/hi-c/publications.html |
| Salameh et al | Peakachu | | K562 | http://promoter.bx.psu.edu/hi-c/publications.html |
| Grubert et al | ChIA-PET | | GM12878 | https://www.ncbi.nlm.nih.gov/pmc/articles/PMC7410831/bin/41586_2020_2151_MOESM5_ESM.xlsx |
| Grubert et al | ChIA-PET | | K562 | https://www.ncbi.nlm.nih.gov/pmc/articles/PMC7410831/bin/41586_2020_2151_MOESM5_ESM.xlsx |

**Table S3.** A complete list of genomic annotations used to build the predictor space for all downstream models. The GRCh37/hg19 human genome assembly was used. "Genomic Class" - broad category of genomic features, "Element" - names of genomic features, "Cell line-Specific Source" - download URL specific to the cell line (not all annotations were provided by the same institutions).

| Tool | Resolution.Bin.size | Total.number.of.called.TADs.chromatin.loops | Total.number.of.unique.TAD.chromatin.loop.boundaries | Total.number.of.genomic.bins | Class.Imbalance |
|---|---|---|---|---|---|
| Arrowhead | 5 kb | 4751 | 9316 | 535363 | 0.02 |
| Arrowhead | 10 kb | 5828 | 10945 | 267682 | 0.04 |
| Arrowhead | 25 kb | 3935 | 7015 | 107073 | 0.07 |
| Arrowhead | 50 kb | 2115 | 3808 | 53537 | 0.07 |
| Arrowhead | 100 kb | 945 | 1759 | 26768 | 0.07 |
| Peakachu | 10 kb | 15651 | 22073 | 267682 | 0.14 |
| Grubert | 5 kb | 12266 | 14325 | 535363 | 0.05 |
